# EHD2 overexpression promotes tumorigenesis and metastasis in triple-negative breast cancer by regulating store-operated calcium entry

**DOI:** 10.1101/2022.06.21.497035

**Authors:** Haitao Luan, Timothy A. Bielecki, Bhopal C. Mohapatra, Namista Islam, Insha Mushtaq, Aaqib M. Bhat, Sameer Mirza, Sukanya Chakraborty, Mohsin Raza, Matthew D. Storck, Michael S. Toss, Jane L. Meza, Wallace B. Thoreson, Donald W. Coulter, Emad A. Rakha, Vimla Band, Hamid Band

## Abstract

With nearly all cancer deaths a result of metastasis, elucidating novel pro-metastatic cellular adaptations could provide new therapeutic targets. Here, we show that overexpression of the EPS15-Homology Domain-containing 2 (EHD2) protein in a large subset of breast cancers (BCs), especially the triple-negative (TNBC) and HER2+ subtypes, correlates with shorter patient survival. The mRNAs for EHD2 and Caveolin-1/2, structural components of caveolae, show co-overexpression across breast tumors, predicting shorter survival in basal-like BC. EHD2 shRNA knockdown and CRISPR-Cas9 knockout of EHD2, together with mouse EHD2 rescue, in TNBC cell line models demonstrate a major positive role of EHD2 in promoting tumorigenesis and metastasis. Mechanistically, we link these roles of EHD2 to store-operated calcium entry (SOCE), with EHD2-dependent stabilization of plasma membrane caveolae ensuring high cell surface expression of the SOCE-linked calcium channel Orai1. The novel EHD2-SOCE oncogenic axis represents a potential therapeutic target in EHD2 and CAV1/2-overexpressing BC.

## Introduction

Breast cancer (BC) remains a major cause of cancer-related deaths, with less than 30% 5-year survival rate in patients with metastatic disease (www.acs.org). Triple-negative BC (TNBC) presents a particularly difficult diagnosis with lack of targeted therapies. A better understanding of tumorigenesis- and metastasis-associated cellular adaptations could open new novel approaches to improve the survival of TN metastatic BC patients.

EPS15-homology (EH) domain-containing (EHD) proteins (EHD1-4) are evolutionarily-conserved lipid membrane-activated ATPases that regulate inward or outward vesicular traffic between the plasma membrane and intracellular organelles by controlling tubulation and scission of trafficking vesicles (1). Unlike other family members, which predominantly localize to endosomal and other intracellular compartments, EHD2 is known to primarily localizes to plasma membrane caveolae to maintain their stable membrane pool (2, 3), suggesting a likely role in caveolae-associated cellular functions. Indeed, caveolae-dependent fatty acid uptake in adipocytes and e-NOS-NO induced small blood vessel relaxation are impaired in EHD2 knockout mice (4, 5). EHD2-dependent stabilization of caveolae was also found to promote the cell surface expression of ATP-sensitive K^+^ channels and protect cardiomyocytes against ischemic injury (6). Caveolae are key to buffering the plasma membrane stress (7) and EHD2 has been shown to positively regulate mechano-transduction through re-localization to the nucleus and by regulation of gene transcription al programs (8).

Recent studies have painted a complex picture of the potential roles of EHD2 in cancer. Reduced EHD2 expression was reported in esophageal, colorectal, breast, and hepatocellular cancers (9–12), with in vitro knockdown or overexpression studies supporting a tumor suppressive role for EHD2. On the contrary, EHD2 overexpression was found as a component of a mesenchymal signature in malignant gliomas with shorter survival, and knockdown analyses showed the EHD2 requirement for cell proliferation, migration, and invasion (13). Higher EHD2 mRNA expression in papillary thyroid carcinomas was associated with extrathyroidal extension, lymph node metastasis, higher risk of recurrence, and presence of BRAF-V600E mutation (14). Studies of clear cell renal cell carcinoma also supported a positive role of EHD2 in tumorigenesis (15). A recent study provided a more mixed picture, with loss of EHD2 expression in TNBC cell lines enhancing their proliferation, migration, and invasion but low levels of EHD2 mRNA in TNBC patient tumors predicting better prognosis (16). Thus, a definitive role of EHD2 in oncogenesis and its mechanisms remain unclear.

Here, our comprehensive expression analyses in BC samples and in vitro and in vivo studies using EHD2 knockdown or knockout approaches in TNBC cell models provide definitive evidence for strong pro-tumorigenic and pro-metastatic role of EHD2. Our studies suggest through a novel pro-oncogenic mechanism of EHD2, namely its requirement for efficient store-operated calcium entry (SOCE), a pathway known to promote tumorigenesis and metastasis in breast and other cancers (17, 18).

## Results

### EHD2 is expressed in basal cells of the mouse mammary gland and in a subset of basal-like breast cancer cell lines

First, we used immunoblotting and immunofluorescence staining of mammary gland tissue from control and *Ehd2*-null mice (generated in the lab; unpublished) to authenticate the specific recognition of EHD2 by an antibody previously validated against ectopic tagged EHD2 (19) (**Supplementary Fig. S1A-B**). High EHD2 expression was seen in mammary adipocytes, consistent with high EHD2 expression in adipose tissues (20). Moderate/high EHD2 staining was seen in the mammary basal/myo-epithelium (smooth muscle actin^+^), but little in the luminal epithelium (cytokeratin 8^+^) (**Supplementary Fig. S1C**). The basal/myoepithelial cell selective localization was confirmed by immunohistochemistry (IHC) (**Supplementary Fig. S1D**). Immunoblotting of basal (EPCAM-low/CD29-high) and luminal (EPCAM-high/CD29-low) mouse mammary epithelial cell-derived organoids further confirmed the basal cell expression of EHD2 (**Supplementary Fig. S1E**). Thus, while mammary adipocytes express the highest levels of EHD2, within the epithelium the basal epithelial cells show selectively express higher EHD2 levels expression.

By immunoblotting, we found EHD2 expression in immortal basal-like mammary epithelial cell lines 76Ntert (hTert-immortalized primary mammary epithelial cell line) (21) and MCF10A, in 2 out of 3 TNBC cell lines, and at lower levels in 3 out of 11 HER2+ cell lines, but in none of the 9 luminal A/B BC cell lines (**Supplementary Fig. S2A**). Immunofluorescence analysis of selected cell lines confirmed the expression pattern seen in immunoblotting and showed exclusive localization of EHD2 to the plasma membrane and cytoplasm (**Supplementary Fig. S2B**). Notably, our cell line results were discordant with reports of comparable EHD2 expression in MCF-7 (luminal), MDA-MB415 (luminal) and MDA-MB-231 (basal) cell lines (11, 22). The pattern of EHD2 protein expression we observed correlated with the EHD2 mRNA expression data in the CCLE database (23) (**Supplementary Fig. S2C**). Furthermore, by relating the CCLE data to the reported BC cell line subtype analysis (24), we found higher EHD2 mRNA expression to be a feature of TNBC cell lines, especially the mesenchymal-type, similar to our western blot results where all 3 EHD2-expressing TNBC cell lines were of the mesenchymal-type (**Supplementary Fig. S2D**). Thus, high EHD2 expression is a feature of normal basal mammary epithelial cells and a subset of the basal-like/triple-negative BC.

### EHD2 overexpression is associated with metastasis and shorter survival in breast cancer

Based on the above findings, we conducted IHC staining of a tissue microarray (TMA) with 840 primary BC samples from a well-annotated patient cohort (25) to assess the expression of EHD2. Given the predominantly cytoplasmic/membrane localization of EHD2 in the mammary gland and BC cell lines, but the reported nuclear localization in cell lines under defined conditions (8, 26), we quantified the IHC signals as cytoplasmic and nuclear (**Fig. 1A**). 759 and 756 cases respectively showed a valid positive/negative cytoplasmic or nuclear signal (**Supplementary Table 1A**). High cytoplasmic and low nuclear EHD2 signals showed a positive association with higher tumor grade, higher mitosis, and lower cyokeratin-5 expression while high nuclear and low cytoplasmic EHD2 signals showed a reverse correlation and was associated with ER/PR/AR-positive and non-TNBC status (**Supplementary Table 1B**). High cytoplasmic EHD2 predicted shorter BC-specific survival, while high nuclear EHD2 showed an opposite correlation (**Fig. 1B**). Across BC subtypes, the high cytoplasmic and nuclear-negative EHD2, which also predicted shorter BC-specific survival (**Supplementary Fig. S2E**), was seen in about half of TNBC and HER2+ samples, and a third of ER+ samples (**Fig. 1C, Supplementary Table 1B**). Analysis of a subset of our patient cohort with data for further subtyping showed a strong skewing of high cytoplasmic EHD2 expression in basal-like TNBC and to some extent the HER2-enriched and luminal B subtypes, while high nuclear EHD2 expression was a feature of luminal A subtype (**Fig. 1D**). Thus, our results indicate that high cytoplasmic EHD2 expression, a localization similar to that observed in normal mammary epithelium and BC cell lines, is a marker of more aggressive BC, contrary to published reports that did not report assess the cytoplasmic/nuclear distribution of EHD2 and suggested its potential tumor suppressor role (11, 16, 22).

**Figure 1.**
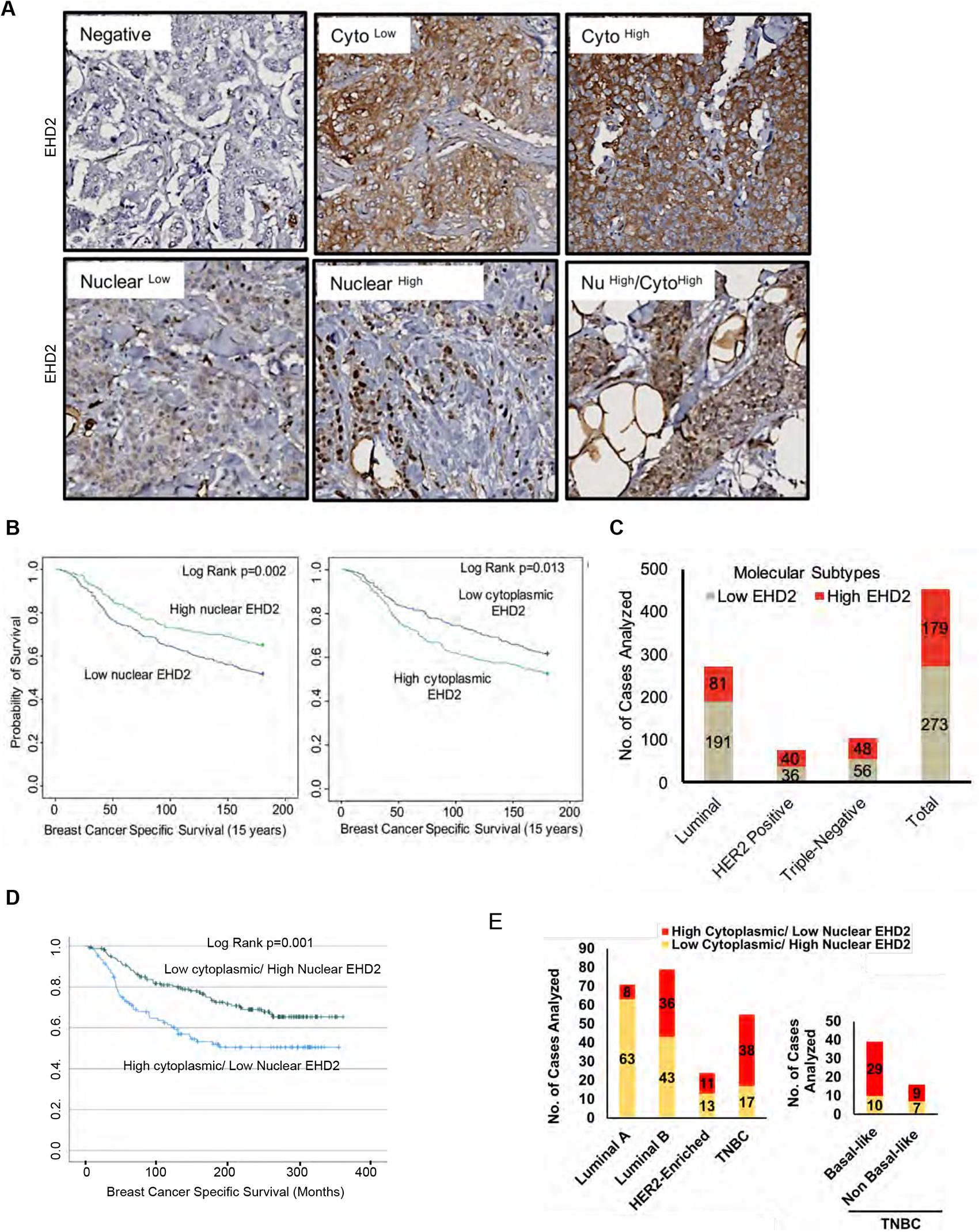
EHD2 is overexpressed in a subset of breast cancer patients and is associated with metastasis and shorter survival. (A) Representative images of negative/low/high cytoplasmic and nuclear EHD2 IHC staining of a breast cancer tumor microarray (840 samples). (B) Kaplan-Meier survival curves correlating positive/high (green) vs. low/negative (blue) nuclear (left panel; N=288 vs. 458) or cytoplasmic (right panel; N=392 vs. 352) EHD2 expression with Breast Cancer Specific Survival (BCSS). (C) Number (Y-axis) of cytoplasmic EHD2-negative/low (gray) and -positive/high samples among ER/PR+, ErbB2+, TN, and all tumors. (D) Left Panel – Kaplan-Meier survival analysis of a subset of patients with molecular subtyping markers available (N=271) comparing high cytoplasmic/low nuclear (blue; N=107 out of 271) vs low cytoplasmic/high nuclear (green; N=164 out of 271). Middle panel – number (Y-axis) of high cytoplasmic/low nuclear EHD2 (red) and low cytoplasmic /high nuclear EHD2 (yellow) cases within the luminal A (ER^+^/PR^+^, HER2^-^ and Ki67 <14%), luminal B (ER^+^/PR^+^ or ^-^ and either HER2^+^ or Ki67 >/= 14% or both), HER2-Enriched (ER^-^, PR^-^ and HER2^+^, regardless of the Ki67) and TNBC (ER, PR and HER2^-^, regardless of the Ki67) BC subtypes. Right panel – number of high cytoplasmic/low nuclear (red) or low cytoplasmic/high nuclear (yellow) EHD2 staining in basal-like (CK5/6 or CK14 or CK17 positive) and non-basal-like (CK5/6, CK14 or CK17 negative) TNBC subtypes.

### EHD2 knockdown or knockout in TNBC cell lines impairs the tumorigenic and pro-metastatic traits

To examine the role of EHD2 expression in BC oncogenesis, we established control or EHD2 shRNA expressing TNBC cell lines, Hs578T, BT549 and MDA-MB-231 (**Fig. 2A**). While EHD2 knockdown (KD) did not affect proliferation in two-dimensional culture on plastic (**Fig. 2B**), it markedly reduced the tumorsphere growth under low attachment (**Fig. 2C**), impaired invasion across Matrigel in trans-well assays (**Fig. 2D**) and markedly reduced the invasive fronts in a Matrigel organoid invasion assay (**Fig. 2E**).

**Figure 2.**
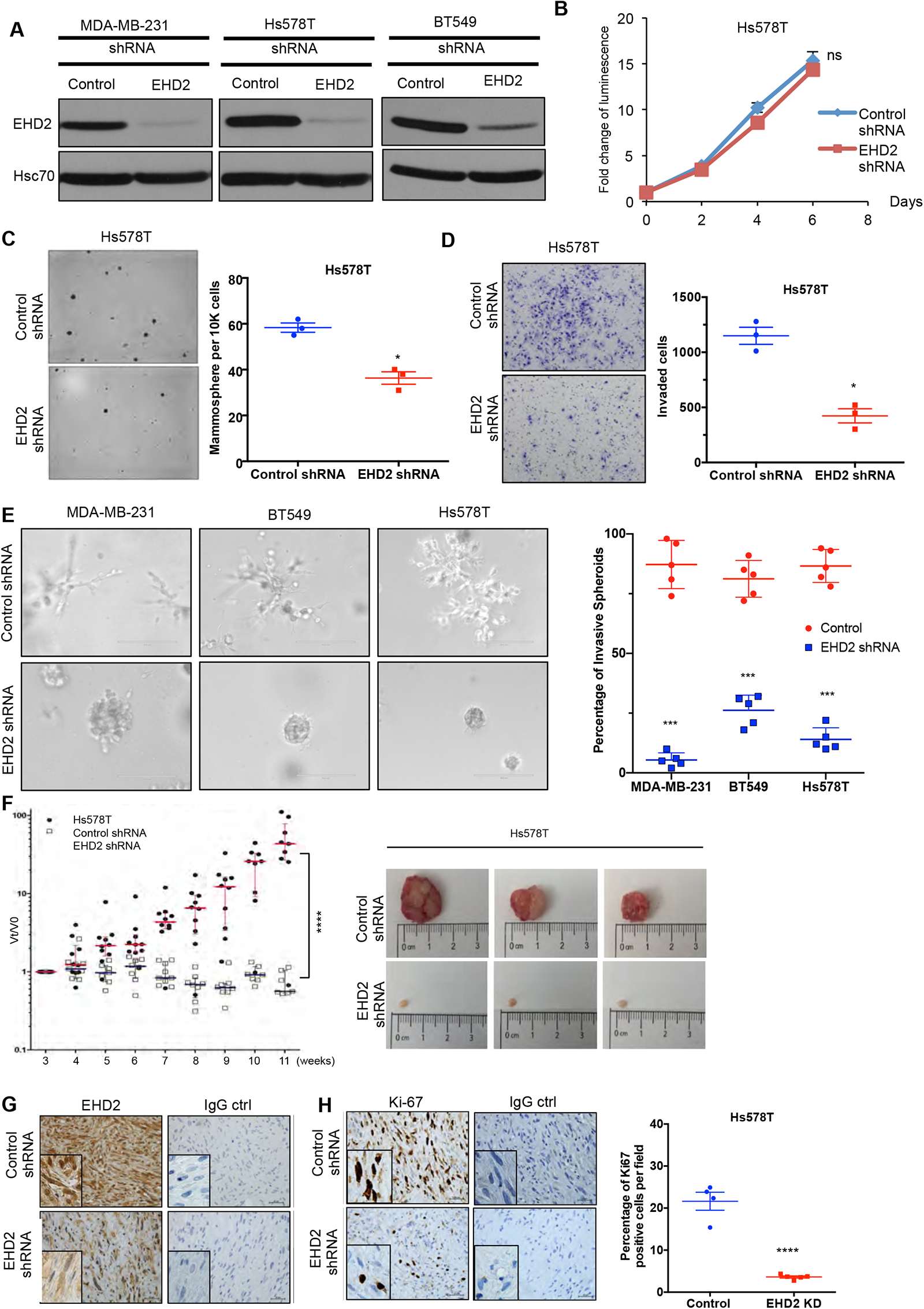
EHD2 knockdown in TNBC cell lines impairs the tumorigenic and pro-metastatic traits. (A) Immunoblot confirmation of shRNA-mediated EHD2 knockdown. (B) Cell Titer-Glo proliferation (2,000 cells/well; 24 replicates each) over time. Mean +/- SEM, n=3, ns, not significant. (C) Tumorsphere formation quantified on day 7. Left, representative images; Right, quantification of tumorspheres/well. Mean +/- SEM, n=3, *p<0.05; **p<0.01. (D) Transwell invasion of cells plated in 0.5% FBS medium towards complete medium assayed after 18h. Left, representative images; Right, quantification of invaded cells (Mean +/- SEM, n=3, *p<0.05). (E) Three-dimensional invasion in Matrigel-grown organoids. 2,000 cells plated per well in 50% Matrigel on top of 100% Matrigel layer in 8-well chamber slides for 7 days before imaging. Left, representative images; right, % spheroids with invasive fronts from over 100 counted per well, n=4, *** p<0.001). (F) Xenograft tumorigenesis. 4-weeks old nude mice orthotopically-injected with 5x10^6^ cells were followed over time. Left, fold change in tumor volume over time for individual mice. Mean (red/blue lines) +/- SEM; ****,p<0.0001 by two-way ANOVA. Right, representative tumors (close to median of groups). (G, H) Representative IHC staining of tumor sections for EHD2 (G) or Ki67 (H), with respective controls. Right, Mean +/- SEM of Ki67+ staining. ****, p<0.0001.

Orthotopically implanted control Hs578T cells expected produced xenograft tumors over time while EHD2 KD cells showed a severe reduction in tumor formation (**Fig. 2F**). Immuno-staining confirmed the EHD2 KD (**Fig. 2G**) and showed marked reduction in proliferation (Ki67^+^) with sparse tumor cells in H&E sections (**Fig. 2H**). EHD2 KD in MDA-MB-231 cells also reduced the xenograft growth and frequency of lung tumor metastasis (**Supplementary Fig. S3**).

Further, CRISPR-Cas9-mediated EHD2 knockout (KO) in TNBC cell lines (**Fig. 3A**), which unlike our observations in *Ehd2*-KO mouse tissues was not associated with any significant changes in EHD1/4 expression (**Supplementary Fig. S4**), significantly impaired their cell migration, invasion (**Fig. 3B-C, Supplementary Fig. S5A-B**) and extracellular matrix (ECM) degradation ability (**Fig. 3D, supplementary Fig. S5C**), another pro-metastatic trait (27). Introduction of mouse EHD2 in MDA-MB-231 EHD2-KO cells, at levels lower than in control cells, significantly rescued the cell migration defect (**Fig. 3E**), demonstrating specificity. Reciprocally, CRISPR activation of endogenous EHD2 in EHD2-nonexpressing MDA-MB-468 TNBC cells (**Fig. 3F**) increased cell migration compared to control cells (**Fig. 3G**). When orthotopically implanted in nude mice, EHD2 KO MDA-MB-231 cells exhibited a marked and significant defect in tumor formation, with a significant rescue upon mouse EHD2 expression (**Fig. 3H**).

**Figure 3.**
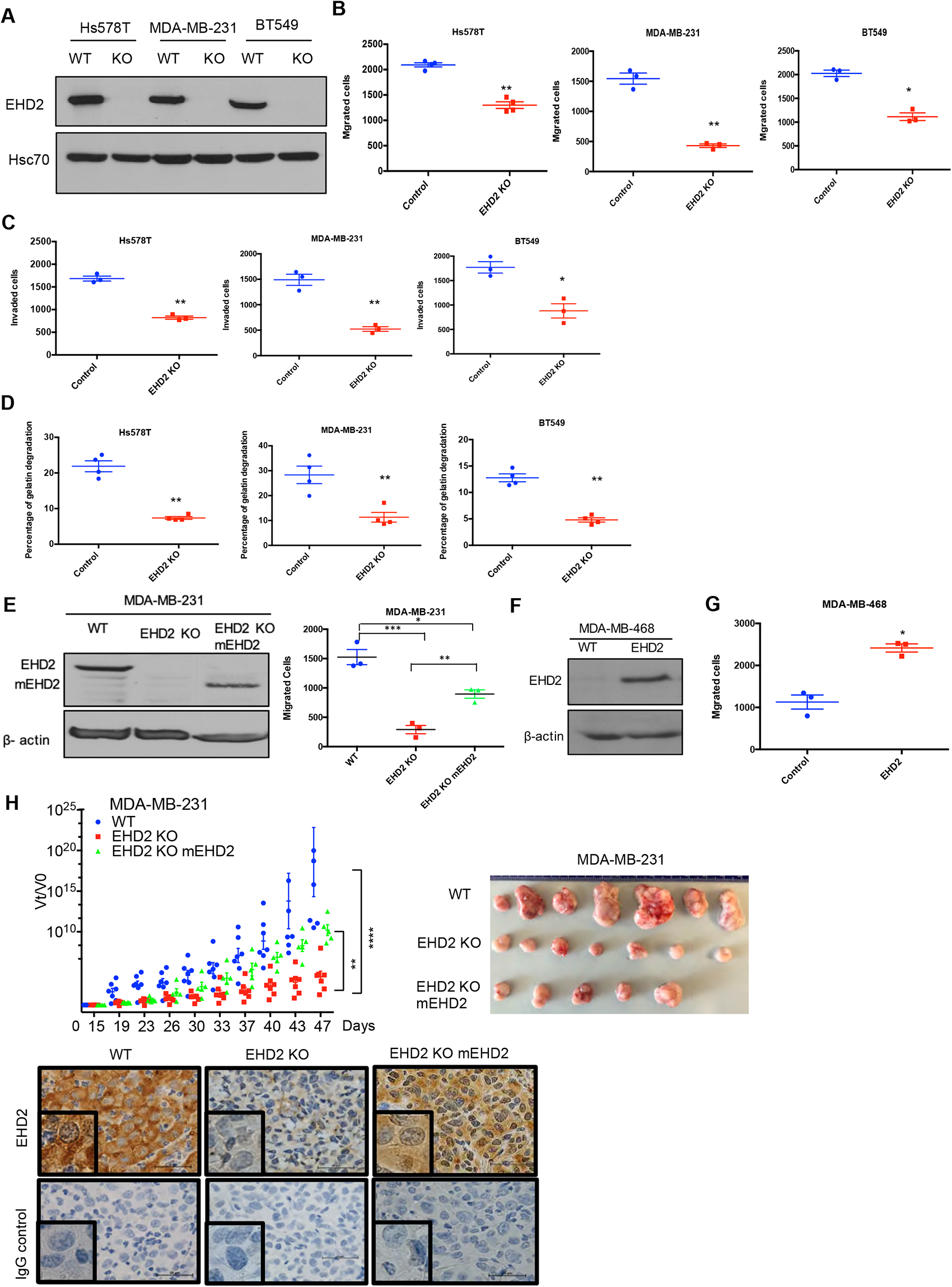
EHD2 knockout in TNBC cell lines impairs the tumorigenic and pro-metastatic traits. Single cell clones of TNBC cell lines serially transduced with Cas9 and control or EHD2 sgRNA lentiviruses were obtained and use as a pool of >3 clones. (A) Immunoblotting of EHD2 expression in KO vs. WT (Cas9) controls. (B) Transwell migration. Data points are independent experiments; Mean +/- SEM of migrated cells (input 10K), **p<0.01, *p<0.05. (C) Transwell invasion across Matrigel. Mean +/- SEM of invaded cells (input 10K), **p<0.01, *p<0.05. (D) Extracellular matrix degradation. Cells plates on Cy5-gelatin and percentage area with matrix degradation quantified after 48h. Mean +/- SEM, **p<0.01. (E) Mouse EHD2 rescue of EHD2 KO MDA-MB-231 cells. Left, immunoblot to show re-expression of mouse EHD2; beta-actin, loading control. Right, rescue of cell migration defect. Mean +/- SEM, ***p<0.001, **p<0.01, *p<0.05. (F-G) CRISPRa induction of endogenous EHD2 expression in EHD2-negative MDA-MB-468 cell line (F) and increase in migration (G). Mean +/- SEM, *p<0.05. (H) Impairment of tumorigenesis by EHD2 KO and rescue by mouse EHD2 reconstitution. Left, groups of 8 nude mice orthotopically implanted with 3x10^6^ cells and tumors analyzed as in Fig. 2F: ****p<0.0001, **p=0.001. Right, Representative tumor images. Bottom, representative tumor sections stained for EHD2 and control.

To directly assess the role of EHD2 in metastasis, luciferase-expressing control and KO MDA-MB-231 cells were intravenously injected into nude mice. Luminescence bioimaging showed time-dependent lung metastatic growth of control cells but no growth (or a reduction in signals) with EHD2 KO cells (**Fig. 4A-C**). These findings were confirmed by assessment of lung metastatic nodules at necropsy (**Fig. 4D**). H&E and human CK18 staining confirmed the metastatic growths, and EHD2 expression pattern was confirmed by IHC (**Fig. 4E**). Collectively, our analyses definitively demonstrate a positive role of EHD2 in tumorigenic and pro-metastatic behavior in TNBC.

**Figure 4:**
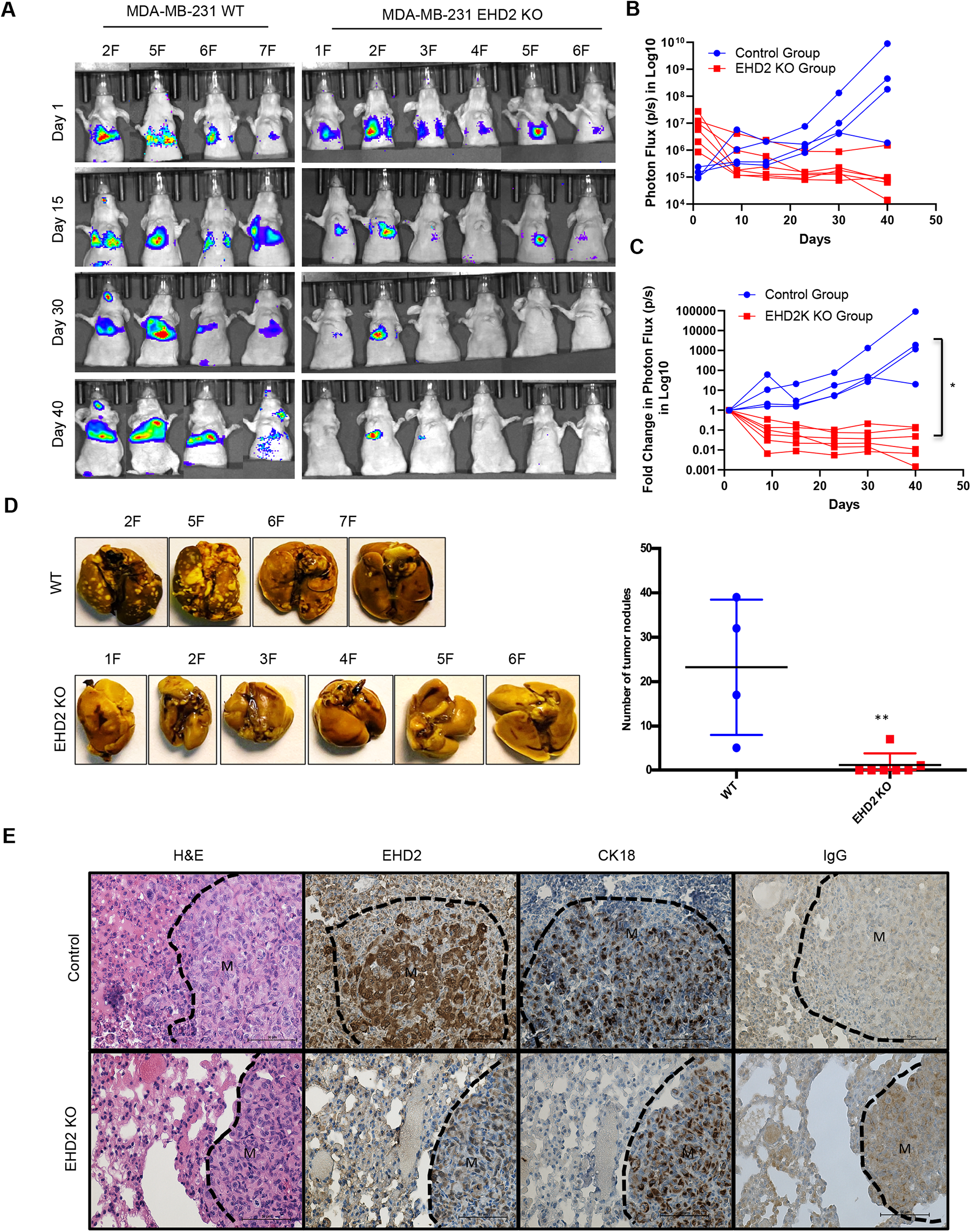
EHD2 KO impairs the ability of TNBC cells to form lung metastases. WT control and EHD2 KO MDA-MB-231 cells were engineered with tdTomato-luciferase and 10^6^ cells of each injected intravenously into groups of 7 nude mice. Lung metastases were monitored by bioluminescence imaging (shown in A). (B-C) Bioluminescence signals over time (Control, blue; KO, red) are shown as either untransformed photon flux values (B) or log fold-change in photon flux relative to day 0 (C). Two-way ANOVA showed the differences between Control and KO groups to be significant (*p<0.05). (D) Left panel, images of lungs harvested at necropsy show nearly complete absence of metastatic nodules in lungs of mice injected with EHD2 KO cells. Right panel, quantification of tumor nodules in the lungs, **, p<0.01. (E) Representative H&E (first panels), EHD2 (second panels), CK18 (third panels) and control IgG staining of metastatic lung tissue sections from control (upper) and EHD2 KO cell injected mice. Note the retention of normal lung tissue in EHD2 KO cell injected mouse lung, and absence of EHD2 expression in KO nodules (labeled M). CK18 demarcates the human tumor cell area.

### EHD2 and CAV1/2 are co-overexpressed in basal-like breast cancer and loss of EHD2 reduces the cell surface caveolae

EHD2 localizes to and is required for the stability of the cell surface caveolae (2, 3, 8, 28). The bc-GenExMiner analysis of 5,277 BC samples (29) demonstrated tight co-expression of EHD2 with the structural components of caveolae, CAV1 and CAV2 in TNBC samples (**Fig. 5A, Supplementary Fig. S6A**). By KM Plotter analysis, combined EHD2-, CAV1- and CAV2-high basal (PAM50-based) but not all BC patients showed significantly shorter distal metastasis-free survival (**Fig. 5B, Supplementary Fig. S6B**). Immunoblotting demonstrated concordant EHD2 and CAV1 expression in mammary epithelial and BC cell lines (**Fig. 5C**). Immunofluorescence analysis using structured illumination microscopy (SIM) demonstrated a high degree of colocalization between EHD2 and CAV1 in TNBC cell lines (**Fig. 5D**). Total internal reflection fluorescence (TIRF) microscopy analysis showed a significant reduction of cell surface associated CAV1-GFP puncta, representing cell surface caveolae, in EHD2 KO compared to control Hs578T cells (**Fig. 5E**), consistent with the reported electron microcopy-based high cell surface caveolae density on Hs578T compared to a lower density on the EHD2-non-expressing MDA-MB436 cells (8). CRISPR KO of CAV1 (**Fig. 5F**) led to a significant impairment of cell migration like that with EHD2 KO (**Fig. 5G**). These results support the conclusion that EHD2-dependent maintenance of cell surface caveolae is linked to its promotion of tumorigenic and pro-metastatic traits, although the potential role(s) of EHD2 and caveolae in other subcellular locations cannot be excluded at present.

**Figure 5.**
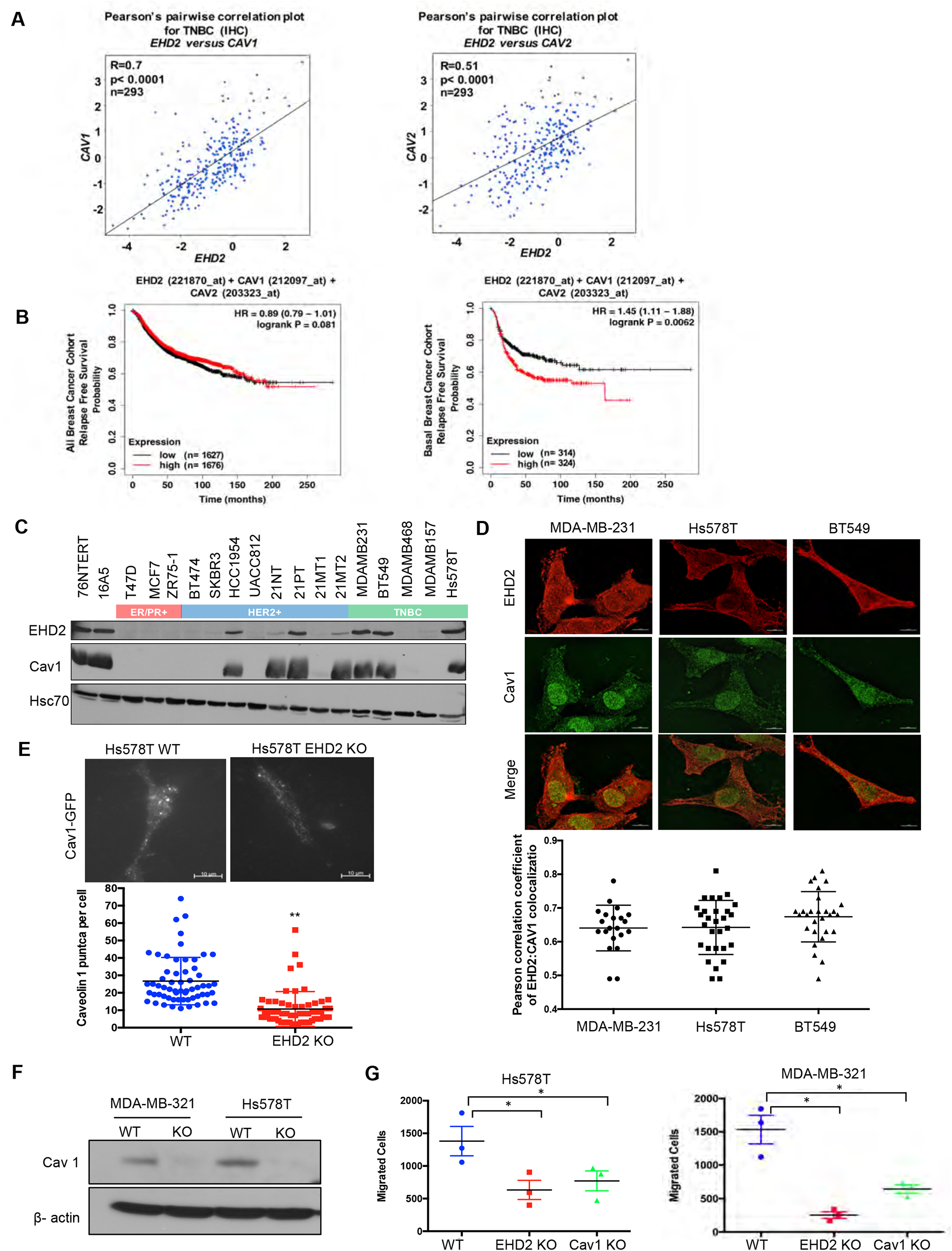
EHD2 and Caveolin-1/2 are co-overexpressed in breast cancers and EHD2 regulates cell surface caveolae. (A) Pearson’s correlation plots of EHD2/CAV1 and EHD2/CAV2 expression in TNBC (IHC-based) subsets of TCGA and SCAN-B RNAseq datasets analyzed on bc-GenExMiner v4.5 platform. Indicated: n, number of samples; R, correlation coefficients; significance. (B) KM plotter analysis of EHD2, CAV1 and CAV2 overexpression correlation with relapse-free survival (RFS) for upper vs. lower quartiles in basal-like breast cancer (PAM50-based) cohorts of TCGA, GEO and GEA datasets. Probe sets used: EHD2 (221870_at), CAV1 (212097_at) and CAV2 (203323_at). Analysis of all samples combined found no survival differences (lower panel). (C) Immunoblot analysis of coordinate EHD2 and CAV1 expression in immortal mammary epithelial cells and breast cancer cell lines. (D) SIM images demonstrated colocalization of EHD2 (red) and caveolin-1 (green) in TNBC cell lines; scale bar, 100 μm. Top, representative SIM images; Bottom, Pearson’s Coefficient of Colocalization between EHD2 and CAV1 in TNBC cells from three independent experiments. (E) TIRF analysis of fluorescent CAV1 puncta to quantify cell surface caveolae pool. Top, representative TIRF images. Bottom, quantification of CAV1 puncta. Mean +/- SEM of puncta per cell pooled from 3 independent experiments; **p<0.01. (F) Immunoblot confirmation of CRISPR-Cas9 CAV1 KO in TNBC cell lines. (G) Impact of CAV1 KO on Transwell migration. Mean +/- SEM number of migrated cells (input 10K) per Transwell (n=3, *p<0.05).

### EHD2 promotes pro-metastatic traits in TNBC cells by upregulating store-operated calcium entry

The I impact of EHD2 depletion on multiple oncogenic traits and its regulation of plasma membrane caveolae suggested the role for a caveolae-linked signaling machinery. We investigated the linkage of EHD2 to store-operated calcium entry (SOCE) (30), a pathway that operates at caveolae (31, 32) and is a well-established pro-metastatic signaling pathway in TNBC and other cancers (17, 18). The SOCE is mediated by translocation of the endoplasmic reticulum (ER) Ca^2+^ sensor to ER-plasma membrane contact sites upon ER Ca^2+^ depletion which permits its binding to and activation of the Orai1 membrane Ca^2+^ channel to promote Ca^2+^ entry for Ca^2+^-dependent signaling and refilling of the ER (30).

To examine if EHD2 regulates SOCE in TNBC cells, calcium-sensitive fluorescent dye (Fluo4 AM)-loaded cells in Ca^2+^ free medium were treated with thapsigargin (Tg), an inhibitor of the ER-localized Sarco-Endoplasmic Reticulum Ca^2+^ ATPase 2 (SERCA-2) (33). Expectedly, control Hs578T or BT549 TNBC cells exhibited a robust rise in cytoplasmic Ca^2+^ that declined over time (**Fig. 6A-B**), reflecting the release of ER Ca^2+^ (34). Subsequent addition of Ca^2+^ in the medium induced a rapid increase in cytoplasmic Ca^2+^, indicating the SOCE (**Fig. 6A-B**) (34). Pre-treatment with the SOCE inhibitor SKF96365 (18) markedly reduced the initial Ca^2+^ flux and nearly abrogated the SOCE (**Fig. 6C**. EHD2 KO cells demonstrated a marked defect in both the initial Tg-induced rise in cytoplasmic Ca^2+^ and the subsequent SOCE (**Fig. 6A-B**)), consistent with the established role of SOCE in intracellular Ca2+ store filling (the source of the initial Ca2+ release upon Tg treatment) besides Ca2+-dependent signaling (30). Defective SOCE was also seen in EHD2 KO Hs578T cells using another SERCA inhibitor cyclopiazonic acid (CPA) (35) (**Fig. 6D**). In a genetic approach, we showed that EHD2 KO Hs578T cells stably expressing a GFP-based reporter of cytoplasmic Ca^2+^, GCaMP6s, (36), exhibited defective Tg-induced SOCE (**Fig. 6E**, **Supplementary Fig. S7A**). Further, stable expression of GCaMP6s-CAAX, a plasma membrane-targeted fluorescent reporter of Ca^2+^ levels (37), which only detects the SOCE phase upon Tg treatment directly established the defective SOCE in EHD2-KO Hs578T cells (**Fig. 6F**, **Supplementary Fig. S7B**). In a reciprocal experiment, CRISPRa-induced endogenous EHD2 expression in EHD2-negative MDA-MB468 cells led to a marked increase in Tg-induced SOCE (**Fig. 6G**). Consistent with the role of caveolae, a marked defect in SOCE was observed in CAV1-KO TNBC cell lines (**Fig. 6H-I**).

**Figure 6.**
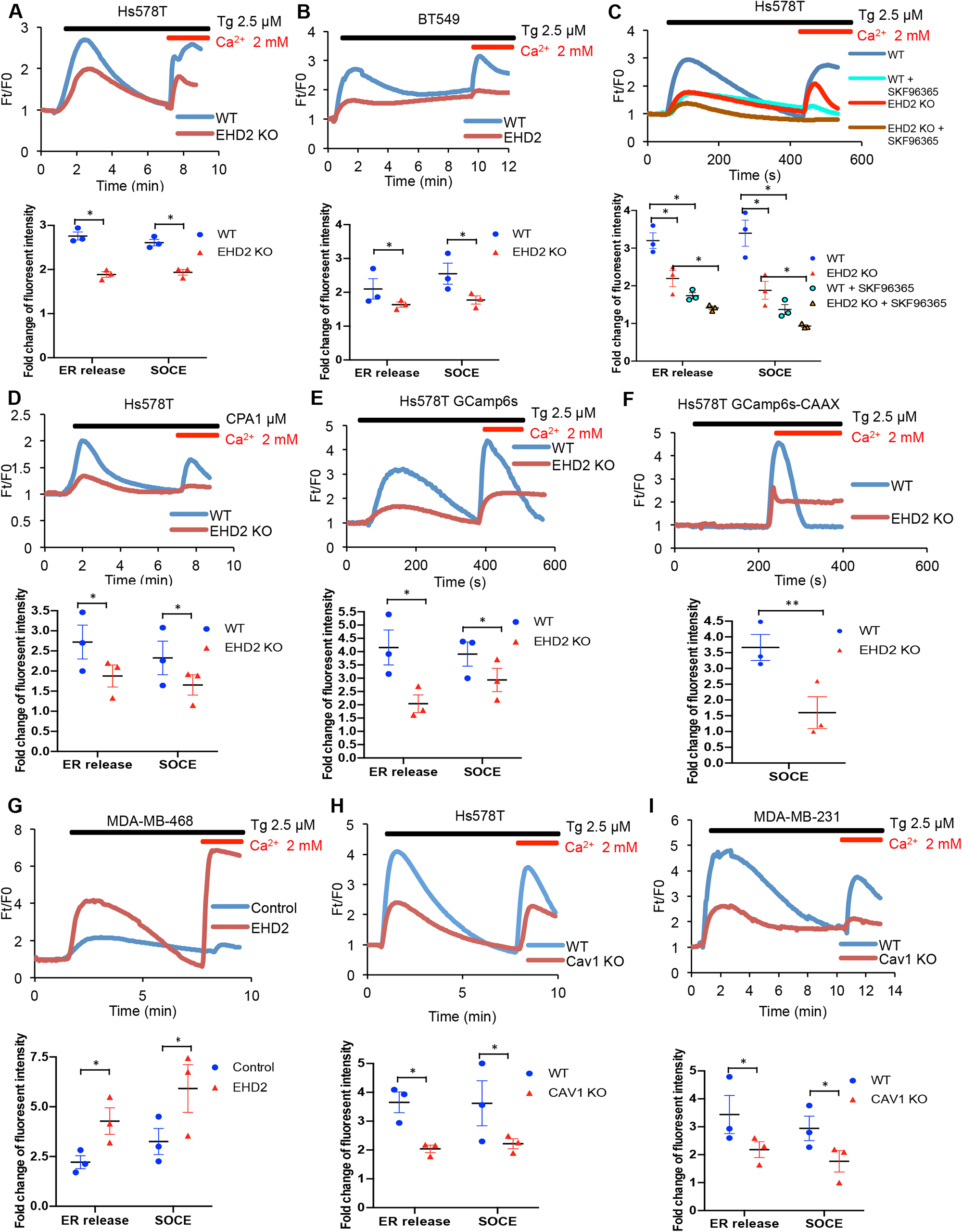
EHD2 promotes store-operated calcium entry (SOCE) in TNBC cell lines. (A-B) Thapsigargin (Tg; 2.5 μM)-induced increase in cytoplasmic Ca^2+^ (initial rise in no extracellular Ca^2+^) and SOCE (second peak after adding 2 mM Ca2+) in Fluo 4 AM-loaded WT/KO Hs578Tin Fluo 4 AM-loaded WT/KO Hs578T (A) or BT549 (B) cell lines measured by live-cell confocal microscopy. (C) Impact of SOCE inhibitor SKF96365 (10 LM) on Tg (2.5 μM)-induced Ca^2+^ fluxes measured as in A. (D) Defective Tg-induced Ca^2+^ fluxes demonstrated using cyclopiazonic acid (CPA; 1-M). (E-F) Tg (2.5 μM)-induced Ca^2+^ fluxes measured by confocal imaging of stably-expressed genetic cytoplasmic Ca^2+^ sensors: cytoplasmic sensor GCaMP6s (E) and plasma membrane-localized sensor GCaMP6s-CAAX (F). (G) Tg (2.5 μM)-induced Ca^2+^ fluxes in Fluo4 AM-loaded control MDA-MB468 (EHD2-negative) vs its CRISPRa derivative (EHD2-expressing). (H-I) Tg (2.5 μM)-induced Ca^2+^ fluxes in Fluo4 AM-loaded control MDA-MB468 (EHD2-negative) vs its CRISPRa derivative (EHD2-expressing). (H-I) Tg (2.5 μM)-induced Ca^2+^ fluxes in Fluo4 AM-loaded control and CAV1 KO TNBC lines. Mean +/- SEM of peak fluorescence intensity (n=3, *p<0.05) is shown below all panels.

Next, we transiently transfected the CFP-tagged STIM1 in control or EHD2 KO Hs578T cells and quantified the number of fluorescent STIM1 puncta at the cell surface, a measure of STIM1-Orai1 interaction, using TIRF microscopy (38). Tg treatment failed to increase the STIM1 puncta in EHD2 KO cells (**Fig. 7A**). This defect was not a result of reduced levels of total STIM1 and Orai1 proteins (**Fig. 7B**). Given the known localization of Orai1 in caveolae (32, 39), we assessed the impact of EHD2 KO on Orai1 cell surface levels. We used an anti-Orai1 antibody authenticated against control or Orai1 knockdown TNBC cell lines (**Supplementary Fig. S8**) to immunoprecipitate Orai1 from surface biotin-labeled control and EHD2 KO MDA-MB-231 or Hs578T cells and confirmed the comparable immunoprecipitation of total Orai1 in WT vs. KO cells (**Fig. 7C, upper lower panels**). In contrast, streptavidin blotting revealed a marked reduction in biotinylated (cell surface) Orai1 signals in EHD2 KO cells (**Fig. 7C, lower upper panels**). Further linking the SOCE to EHD2-dependent pro-metastatic traits, overexpression of CFP-STIM1 in EHD2-KO Hs578T cells (**Fig. 7D**) partially rescued the SOCE defect (**Fig. 7E**) and the defective cell migration (**Fig. 7F**). In a complementary approach, the tool SOCE inhibitor SKF96365 and a recently identified inhibitor CM4620, which (as Auxora^TM^, CalciMedica) has progressed to phase 3 clinical trials in acute inflammatory disease conditions (40), impaired the SOCE in wildtype TNBC cells and further reduced the residual SOCE in EHD2-KO cells (**Fig. 6C**). These SOCE inhibitors significantly impaired the wildtype TNBC cell migration, and further reduced the migration of EHD2 KO cells, albeit it the latter was not statistically significant (**Fig. 8A**). Thus, a major proportion of the SOCE in TNBC cell lines is dependent on EHD2 and is inhibitable with available SOCE inhibitors. Accordingly, we show that SKF96365 treatment significantly reduced the control TNBC xenograft tumor growth (**Fig. 8B**); we were unable to test the SOCE inhibitor against EHD2 KO TNBC xenografts as these did not grow sufficiently to test the inhibitor impact of inhibition. Collectively, these findings support our conclusion that EHD2, by stabilizing caveolae, facilitates the SOCE to promote downstream pro-oncogenic traits in TNBC cells.

**Figure 7.**
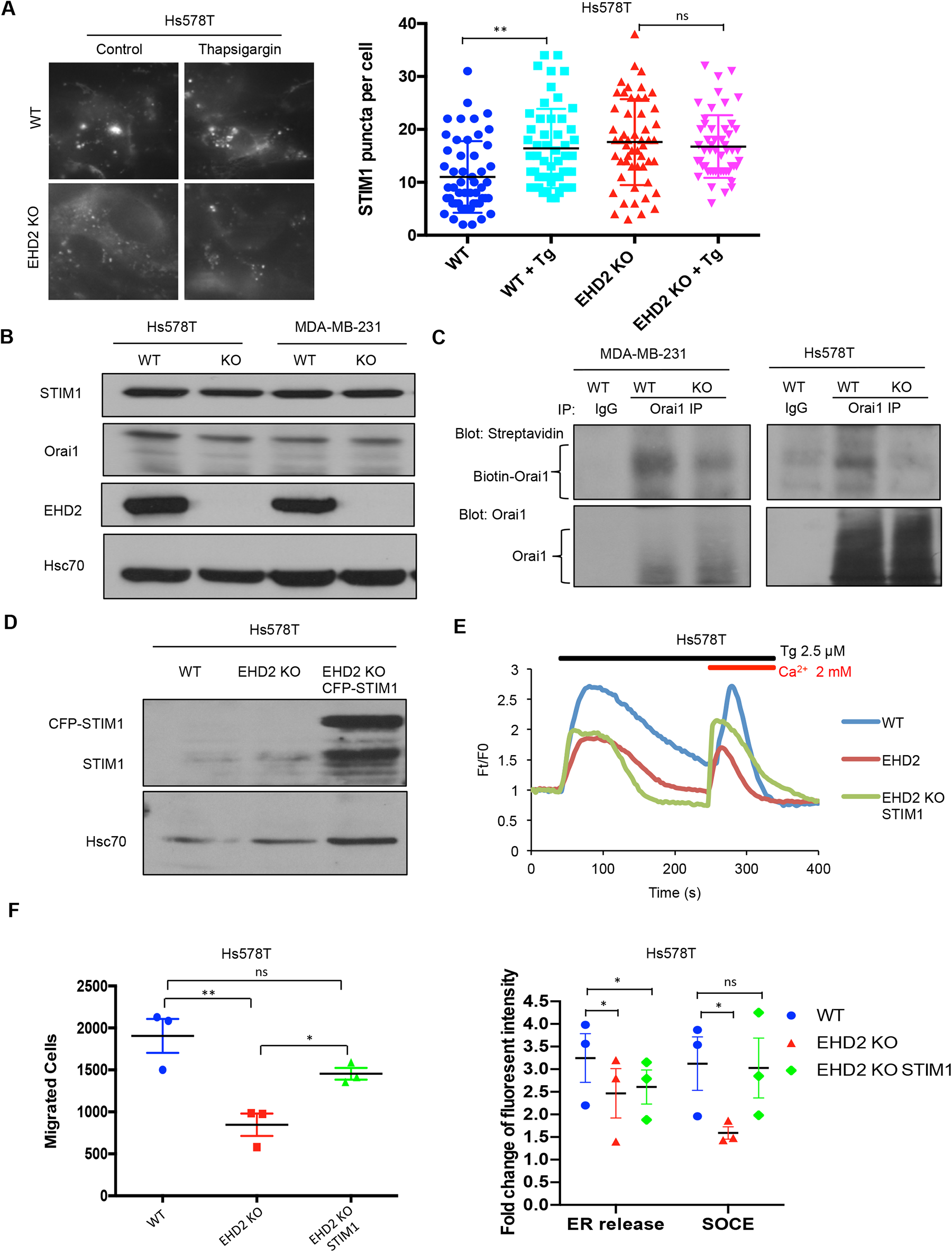
EHD2 regulates SOCE through STIM1-Orai1. (A) CFP-STIM1-trasnfected cells were analyzed for plasma membrane proximal fluorescent puncta by TIRF microscopy, without (control) or with thapsigargin treatment (2.5 L Right, Mean +/- SEM of STIM1 puncta/cell, ** p<0.01. (B) Immunoblotting to show comparable total STIM1 and Orai1 levels in control vs EHD2 KO TNBC lines; Hsc70, loading control. (C) Reduced cell surface levels of Orai1 in EHD2 KO cells. Live cell surface biotinylated cell Orai-1 immunoprecipitates blotted with Streptavidin (top) and Orai1 (bottom). (D) Anti-STIM1 immunoblotting to show stable overexpression of STIM1-CFP in EHD2 KO Hs578T cells. (E) Partial rescue of SOCE by ectopic CFP-STIM1 overexpression analyzed upon thapsigargin (Tg; 2.5 μM) treatment of Fluo 4 AM-loaded cells. Bottom, Mean +/- SEM of peak fluorescence, N=3; *p<0.05. (F) Partial rescue of Transwell cell migration defect by CFP-STIM1 overexpression in EHD2 KO cells. Mean +/- SEM of migrated cells (input 10K); n=3; *P<0.05.

**Figure 8.**
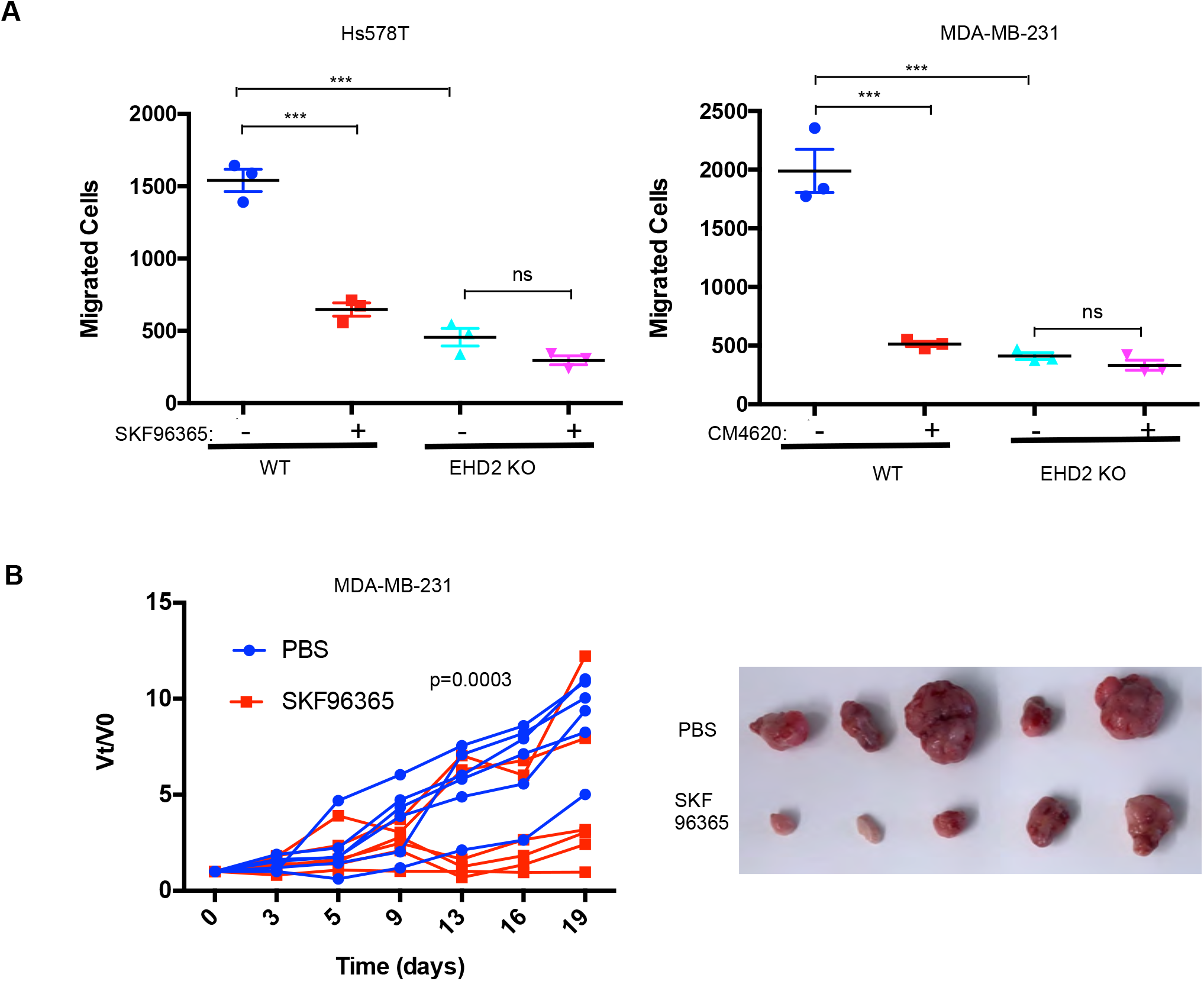
EHD2 expression determines the relative functional impact of SOCE inhibition in TNBC cells. (A) Impact of SOCE inhibitors SKF96365 (10 L M) Transwell migration of Control vs EHD2 KO TNBC cell lines. Mean +/- SEM of n=3; ***P<0.001. (B) SOCE inhibition reduces TNBC tumorigenesis. Nude mice (8/group) bearing orthotopic MDA-MB-231 (3x10^6^) cell tumors (average 4-5 mm in diameter) were administered 10 mg/kg SKF96365 (in PBS) or PBS intraperitoneally and change in tumor volumes (V_t_/V_0_) monitored over time and differences analyzed by two-way ANOVA. Right, representative tumor images.

## Discussion

Elucidating novel tumorigenesis- and metastasis-associated cellular adaptations could dictate new therapeutic options in BC. Here, we use TNBC cell models to elucidate a novel signaling axis linking EHD2 overexpression in BC to store-operated calcium entry (SOCE), a known pro-oncogenic and pro-metastatic pathway. Our studies support the potential for targeting the SOCE pathway in EHD2-overexpressing TNBC and other BC subtypes.

Our IHC analyses demonstrated high cytoplasmic EHD2 expression in a substantial proportion of breast tumors, associated with shorter BC-specific patient survival (**Fig. 1B**) and higher tumor grade (**Supplementary Table 1**). A higher proportion of TNBC, luminal B and HER2+ patients exhibited high cytoplasmic EHD2 (**Fig. 1D, Supplementary Table 1**). Our results differ from reports that reduction in EHD2 expression correlates with tumor progression in BC (11, 22). Notably, a recent study, while it reported the depletion of EHD2 to an increase the oncogenic traits of BC cell lines in vitro, found low EHD2 expression in breast tumors to specify good prognosis and better chemotherapy response (16). Several factors could account for the discordance, including the lack of antibody validation of antibodies in prior studies, the high EHD2 expression in normal mammary adipocytes (**Supplementary Fig. S1A-B**), resulting in apparent reduction in EHD2 expression in tumor tissue using western blotting (11, 16, 22), and the possibility that EHD2 signals in prior studies represented nuclear EHD2, which we find is associated with positive prognostic factors (**Fig. 1B, Supplementary Table 1**). Previous cell line-based studies have shown that EHD2 shuttles between the nucleus and cytoplasm based on nuclear import and exit signals, the latter dependent on its SUMOylation, and that nuclear EHD2 repressed transcription from fused transactivation domains (26). Another study found that application of mechanical stress to cells led to EHD2 release from caveolae, its SUMOylation and nuclear translocation where it regulated transcription of several genes including caveolar components, in a potential positive feedback mechanism stimuli were reported to promote the nuclear translocation of EHD2 (8). However, but any no roles have thus far been assigned to link of the nuclear/cytoplasmic partitioning of EHD2 to in oncogenesis is currently unknown. In view of the opposite prognostic significance of nuclear vs. non-nuclear EHD2 in our analyses of BC patient samples, it could be speculated that nuclear translocation may sequester EHD2 to inhibit its plasma membrane-associated pro-oncogenic role in SOCE, but rigorous studies will be needed to test this suggestion.

Analysis of mRNA expression in public BC databases (**Fig. 5A**) and of protein levels in BC cell lines (**Fig. 5C**) demonstrated high degree of EHD2 co-expression with caveolin-1/2, the structural elements of caveolae. This was noteworthy since EHD2 is known to regulate the stability of caveolae (2, 3). Significantly, EHD2 and/or caveolin-1/2 mRNA overexpression predicted shorter patient survival specifically in the PAM50-defined basal BC (**Fig. 5B**), consistent with the predominant basal (myoepithelial) cell expression of EHD2 in mouse mammary epithelium (**Supplementary Fig. S1B, 1D**). Additional analyses of publicly-available cell line mRNA data in CCLE (**Supplementary Fig. S2C**) and of a subset of BC samples analyzed for EHD2 expression (**Supplementary Fig. S2A**), further reinforce the conclusion that Thus, high cytoplasmic EHD2 expression is a feature of BC with basal-like/triple-negative features.

Multi-pronged approaches using shRNA knockdown and CRISPR-Cas9 KO of EHD2 in TNBC cell models together with mouse rescue of EHD2-KO demonstrated that EHD2 is required for tumorigenesis and metastasis. We show that in vitro tumor cell growth under stringent conditions (tumorspheres) (**Fig. 2C**) and pro-metastatic traits of cell migration, invasion, and ECM degradation (**Fig. 2D, 3B, 3D**) are EHD2-dependent. In vivo, loss of EHD2 markedly impaired the orthotopic TNBC xenograft formation and metastasis (**Fig. 2F**; **Fig. 3H**), and tumor growth was rescued by exogenous mouse EHD2 (**Fig. 3H**). Notably, intravenous injections demonstrated the inability of EHD2-KO TNBC cells to form lung metastases (**Fig. 4A-C**). Although further studies are needed to fully dissect the steps at which EHD2 is critical in the metastatic process, collectively, our analyses conclusively demonstrate that extranuclear EHD2 overexpression in BC cells represents a key pro-tumorigenic and pro-metastatic adaptation.

Mechanistically, we link the EHD2 overexpression in BC cells to regulation of caveolae, whose stability is known to be controlled by EHD2 (2, 3, 20). This includes the strong EHD2 co-localization with CAV1/2 (**Fig. 5D**), reduction in cell surface caveolae density using TIRF microscopy upon EHD2 KO (**Fig. 5E**), and inhibition of TNBC cell migration upon CAV1 KO (**Fig. 5G**), consistent with the previously documented pro-tumorigenic roles of CAV1 in TNBC (41, 42). However, since our localization analyses show both EHD2 and CAV1 to also localize intracellularly besides at the plasma membrane (**Fig. 5D**), and prior studies have identified pro-oncogenic roles for CAV1 localized in various intracellular locations (43), further studies will be needed to determine if the role of EHD2 we define here is exclusively related to regulation of the plasma membrane caveolae or might involve CAV1 in other compartments as well.

Caveolae serve as hubs for signaling (44). Among these, the SOCE pathway stood out as it is known to regulate multiple tumorigenic and pro-metastatic traits in TNBC (31, 32), as with EHD2 depletion. Also, EHD2 interacts with Ca^2+^-binding proteins such as Ferlins (45) that are involved in Ca^2+^-dependent membrane repair and EHD2 was found to accumulate at sites of membrane repair in skeletal muscle models (46, 47). Indeed, our extensive analyses demonstrate that EHD2 is a major positive regulator of SOCE in TNBC cell models. This includes analyses of fluorescent dye-labeled cells and two distinct agents (thapsigargin or CPA) to release ER Ca^2+^ as stimuli (**Fig. 6A-B**), and independent validation using cytoplasm- or plasma membrane-localized genetic reporters of Ca^2+^ (**Fig. 6E-F**, **Supplementary Fig. S7A-B**). Reciprocally, CRISPR-activation of endogenous EHD2 expression in an EHD2-nonexpressing TNBC cell line upregulated SOCE (**Fig. 6G**) and cell migration (**Fig. 3G**). Complementing these, EHD2 KO reduces the STIM1-Orai1 interaction at the ER-plasma membrane contact sites as measured using fluorescent STIM1 (**Fig. 7A**) and overexpression of STIM1 partially rescues the SOCE and cell migration defects in EHD2-KO TNBC (**Fig. 7E-F**). Consistent with the established role of SOCE in the intracellular Ca2+ store refilling (30), which then serves as the source of the initial Ca2+ release upon appropriate stimuli (Tg or CPA in our studies), loss of EHD2 or CAV1 expression or treatment with SOCE inhibitors led to an impairment of the SOCE as well as the initial Ca2+ peak (**Fig. 6A-E**; **Fig. 6H-I**). Reciprocally, CRISPRa upregulation of EHD2 elevated both phases of Ca2+ flux (**Fig. 6G**).

Orai1 is a major STIM1-interacting caveolae-resident SOCE channel (32, 39). Indeed, our cell surface biotinylation studies demonstrated that EHD2 KO specifically reduces the cell surface Orai1 levels (**Fig. 7C**). Thus, our findings support a model whereby EHD2-dependent stabilization of cell surface caveolae ensures high cell surface levels of Orai1 to enable robust SOCE in TNBCs, which in turn promotes pro-tumorigenic and pro-metastatic behaviors of tumor cells (17, 18). Consistent with this model, EHD2 deficiency reduced the cell surface levels of caveolae-associated ATP-sensitive K^+^ channels (6). However, further genetic studies to perturb Orai1 and its family members as well as other potential mediators of SOCE will be necessary to fully establish a causal role of the SOCE pathway in EHD2-dependent oncogenesis.

Finally, consistent with prior studies (18), chemical inhibition of SOCE markedly impaired the pro-metastatic traits of EHD2-overexpressing TNBCs, with a smaller impact on EHD2 KO cell lines (**Fig. 8A**) and impaired the TNBC metastatic growth in vivo (**Fig. 8B**). However, since SKF96365 targets additional Ca2+ channels besides Orai1 (48, 49), further studies using more selective inhibitors together with genetic analyses will be needed. Together, our studies support the idea that EHD2-overexpressing subsets of TNBC and other BC subtypes may be selectively amenable to SOCE targeting, with EHD2 and CAV1/2 overexpression as predictors of response.

## Materials and methods

### Cell lines and Medium

All breast cancer cell lines were obtained from ATCC and cultured in -MEM medium with 5% fetal bovine serum, 10 mM HEPES, 1 mM each of sodium α pyruvate, nonessential amino acids, and L-glutamine, 50 μM 2-ME, and 1% penicillin/streptomycin (Life Technologies, Carlsbad, CA). The TNBC cell lines BT549 and Hs578T were cultured in α-MEM medium supplemented as above and with 1 μ/mL hydrocortisone and 12.5 ng/mL epidermal growth factor (Millipore Sigma, St. Louis, MO). The hTERT-immortalized 76NTERT (21) and spontaneously immortalized MCF10A (50) human mammary epithelial cell lines were maintained in DFCI-1 medium, which contains 12.5 ng/ml EGF. Cell lines were maintained for less than 90 days in continuous culture and were regularly tested for mycoplasma.

### Antibodies and Reagents

Antibodies used for immunoblotting were as follows: HSC70 (# sc-7298) and TRPC1 (# sc-133076) from Santa Cruz Biotechnology; STIM1 (# ab57834) from Abcam; Orai1 (# O8264) and beta-actin (# SAB1305567) from Millipore-Sigma; Caveolain-1 (#610057) from BD Biosciences. In-house generated Protein G-purified rabbit polyclonal rabbit anti-EHD2, antisera has been described previously(19). The horseradish peroxidase (HRP)-conjugated Protein A or HRP-conjugated rabbit anti-mouse secondary antibody for immunoblotting were from Invitrogen. The alpha smooth muscle actin (# ab7817), cytokeratin 8 (#53280), cytokeratin 18 (# 133263), Ki67 (# ab16667) antibodies for immunohistochemistry (IHC) and immunofluorescence (IF) staining were from Abcam. Thapsigargin (# T7459) and Fluo 4AM (# 14201) were from Thermo Fisher Scientific. Cyclopiazonic acid (# C1530) and D-luciferin (#L9504) were from Millipore Sigma. SKF96365 (cat. # S7999), SOCE inhibitor CM4620 (# S6834) from SelleckChem, Matrigel (# 356230) from Corning, and Isoflurane (# 502017) from MWI Animal Health.

### Transfection Reagents and Plasmids

XtremeGENE 9 transfection reagent was from Roche Applied Science (Indianapolis, IN); Commercial pQCXIX-RT3GEP (51) shRNA construct for scrambled shRNA or EHD2-targeted shRNA (Sequence 1, GAAGGCTCGAGAAGGTATATTGCTGTTGACAGTGAGCGCTCACGCTTACATCATCAGCTATAGTGAAGCCACAGATGTATAGCTGATGATGTAAGCGTGAATGCCTACTGCCTCGGACTTCAAGGGGCTAGAATTCGAGCA; Sequence 2, GAAGGCTCGAGAAGGTATATTGCTGTTGACAGTGAGCGCTCCATCCGTCATTCATTCAAATAGTGAAGCCACAGATGTATTTGAATGAATGACGGATGGATTGCCTACTGCCTCGGACTTCAAGGGGCTAGAATTCGAGCA) were custom-made through Mirimus (Brooklyn, NY). Human STIM1-CFP plasmid (52) was from Addgene (#19755). CAV1-mEGFP plasmid (53) was from Addgene (#27704). Lentiviral Mouse EHD2 vector (pLenti-GIII-CMV-GFP-2A-puro, cat. # 190520640395) was from Applied Biological Materials (Richmond, BC, Canada). Luciferase/tdTomatao reporter was engineered using the MuLE system kit from Addgene (Cat. # 1000000060) (54).

### Generation of shRNA knockdown cell lines

Retroviral production and infection was carried out as described previously (55). Briefly, for retroviral expression of control and EHD2 shRNA, HEK-293T cells were transfected with pQCXIX-RT3GEP vectors harboring control shRNA or EHD2 shRNAs together with packaging vectors (plK was a gift from Dr. David Root, Broad Institute), and supernatants used to infect triple negative breast cancer cell lines followed by selection in puromycin.

### Generation of CRISPR-Cas9 knockout/activation cell lines

Edit-R Lentiviral Cas9 nuclease and Edit-R Lentiviral sgRNA control and EHD2 vectors (# SO-2646983G, Dharmacon) were used in a two-step transduction process to derive CRISPR-Cas9 EHD2 KO cell lines. CAV1 sgRNA CRISPR/Cas9 All-in-One Lentivector (pLenti-U6-sgRNA-SFFV-Cas9-2A-Puro) from Applied Biological Materials (Richmond, BC, Canada) was used to derive CAV1 KO cell lines. For EHD2 amplification, dCas9 Synergistic Activation Mediator Lentivector (pLenti-EF1a-dCas9-SAM, cat. # K015) and EHD2 CRISPRa sgRNA Lentivector (pLenti-U6-sgRNA-PGK-Neo, cat. # K0663271) from Applied Biological Materials (Richmond, BC, Canada) were serially infected into MDA-MB468 cell line. In all cases, clonal derivatives were obtained by limiting dilution and screened for complete KO using western blotting. Unless indicated, 3 or 4 clones (maintained separately) were pooled for experimental analyses.

### Cell Lysates

Cells were either lysed in RIPA (50 mM Tris pH 7.5, 150 mM NaCl, 1% Triton-X-100, 0.05% deoxycholate, 0.1% SDS, 1 mM phenylmethylsulfonyl fluoride (PMSF), 10 mM NaF, and 1 mM sodium orthovanadate) or Triton-X-100 (50 mM Tris pH 7.5, 150 mM NaCl, 0.5% Triton-X-100, 1 mM PMSF, 10 mM NaF, and 1 mM sodium orthovanadate) lysis buffer. Lysates were rocked at 4°C for at least 1 hour, spun in a microfuge at 13,000 rpm for 20 minutes at 4°C and supernatant protein concentration determined using the BCA (Thermo Fisher Scientific, Rockford, IL) or Bradford (Bio-Rad Laboratories, Hercules, CA) assay kits.

### Immunoprecipitation

1 mg lysate protein aliquots were used for immunoprecipitation with optimized amounts of the indicated antibodies and 20 μL of Protein A Sepharose beads (GE Healthcare, Chalfont St. Giles, UK) followed by SDS-Polyacrylamide (Biorad) Gel Electrophoresis (PAGE), Polyvinylidene fluoride (PVDF) membrane (Bio-Rad Laboratories, Hercules, CA) transfer and Western Blotting, as described (56).

### Immunofluorescence microscopy

Immunofluorescence staining was performed as described (57) with minor modifications. Cells cultured on glass coverslips were fixed with 4% PFA/PBS (10 min), blocked with 5% BSA/PBS (60 min), and incubated with primary antibodies in 5% BSA/PBS overnight at 4°C. Coverslips were washed with PBS (3x), incubated with the appropriate fluorochrome-conjugated secondary antibody for 45 min at room temperature (RT), washed and mounted using VECTASHEILD mounting medium (cat. # H-1400, Vector Laboratories). For Structured Illumination Microscopy (SIM), images were acquired using a Zeiss ELYRA PS.1 microscope (Carl Zeiss). The Pearson coefficient of colocalization was determined using the FIJI package of ImageJ (NIH). For tissue staining, rehydrated tissue sections were boiled in antigen unmasking solution (H-3300, Vector Laboratories, Burlingame, CA) in a microwave for 20 min, slides were cooled, washed once in PBS, and blocked in heat-inactivated 10% FBS (SH30910.03, HyClone Laboratories, Logan, UT) for 1h at RT. Primary antibodies diluted in blocking buffer were added overnight at 4°C, slides were washed 3x with PBS followed by incubation with Alexa Fluor 488 or 594-conjugated donkey anti-rabbit or anti-mouse secondary antibodies (Invitrogen, Carlsbad, CA) for 1h at RT in the dark. For negative controls, sections were incubated in the blocking buffer without the primary antibody. Nuclei were visualized with DAPI in antifade mounting medium (ProLong® Gold Antifade mountant, Invitrogen, Carlsbad, CA). Fluorescence images were captured on a Zeiss LSM-710 confocal microscope.

### Proliferation assay

A Cell Titer-Glo assay was performed according to the manufacturer’s specifications (Promega, Madison, WI).

### Extracellular matrix degradation assay

This assay was carried out using QCM™ Gelatin Invadopodia kit (Cat. # ECM670, EMD Millipore, Billerica, MA) according to the manufacturer’s protocol. FITC-labeled gelatin was coated onto glass coverslips and crosslinked with 0.5% glutaraldehyde in PBS for 30 min. Coated coverslips were then washed 3x each with PBS and 50 mM glycine in PBS. Cells were cultured for various time points to allow ECM degradation, seen as focal loss of fluorescent signal (“holes”) in the labeled gelatin layer. The fluorescence intensity was further analyzed using the Image J software (NIH).

### Anchorage-independent growth assay

2,500 cells were seeded in 0.35% soft agar on top of 0.6% soft agar layer in 6-well plates. After two weeks, cells were stained with crystal violet and imaged under a phase contrast microscope. The number of colonies in the plate were enumerated using the Image J software (NIH). All experiments were done in triplicates and repeated 3 times.

### Tumor-sphere assay

Cells were re-suspended in DMEM/F12 media (# SH30023.01, GE Lifesciences) supplemented with 2mM L-glutamine, 100U/ml penicillin/streptomycin, 20ng/ml EGF (sigma), 10 ng/ml FGF (R&D Systems) and 1x B27 supplement (Gibco) and seeded at 100,000 cells/well in poly-HEMA coated 6-well plates. After one week, tumor-spheres were imaged under a phase contrast microscope. Tumorspheres greater than 40 μm in diameter were quantified as previously described (58)using the Image J software (NIH, MD). All experiments were done in triplicates and repeated 3 times.

### Matrigel spheroid assay

Cells were re-suspended as a single cell suspension in media containing 50% Matrigel and seeded at 2,000 cells/well on top of a base layer of Matrigel in an 8-well chamber slide. TNBC cell spheroids were allowed to grow for 7 days and then imaged and quantified. Quantification of invasive spheroids was performed by comparing the number of spheroids with invadopodia to the total number of spheroids. Over 100 total spheroids were counted per well and all experiments were done in triplicates and repeated three times.

### Trans-well migration and invasion assays

For migration assays, the cells were seeded in the top chambers of trans-wells (cat. # 353097, Corning) in serum-free medium. For invasion assay, the cells were seeded Matrigel-precoated top chambers of trans-wells (cat. # 354480, Corning) in serum-free medium. Medium containing 10% FBS in the lower chamber served as a chemoattractant. After 18 h, the cells on the upper side of the membrane were removed by scraping with cotton swabs, and cells on the lower side were fixed with methanol, stained with crystal violet, and counted. Experiments were run in triplicates and repeated three times.

### Orthotopic xenograft tumorigenesis

3 million cells in 0.1 ml 50% Matrigel (BD Biosciences) were implanted in the mammary fat pads of 4-6 weeks old non-pregnant female athymic nude mice (The Jackson Laboratory). Tumor growth was monitored using calipers weekly for up to 10 weeks. Tumor volume was calculated as length x width x depth/2 (56). Mice were euthanized when control tumors reached 2 cm^3^ in volume or showed signs of ill health, as per institutional IACUC guidelines. At the end of the experiment the primary tumor, liver and lung were resected, formalin-fixed and paraffin-embedded for further analyses.

### Analysis of tumor metastasis after tail vein injection of tumor cells

10^6^ control or EHD2 KO MDA-MB-231 cells engineered with tdTomato/luciferase reporter were resuspended in 0.1 ml PBS and injected into the lateral tail-vein of 5 weeks-old non-pregnant female athymic nude mice. For bioluminescent imaging, mice received an intraperitoneal injection of 200 μl D-luciferin (15 mg/ml; cat. # L9504 from Millipore Sigma) 15 min before isoflurane anesthesia and were placed dorso-ventrally in the IVIS™ Imaging System (IVIS 2000). Images were acquired using the IVIS Spectrum CT and analyzed using Living Image 4.4 software (PerkinElmer). Mice were imaged weekly and followed for up to 40 days. At the end of the study, lungs were harvested from euthanized mice, fixed in paraformaldehyde, and embedded in paraffin for histopathological analysis.

### TIRF microscopy

Cells were seeded on 1.78 refractive index glass coverslips and transfected with pGFP-CAV1 (for CAV1 puncta) or STIM1-CFP (for STIM1 puncta). Cells were treated with or without thapsigargin (2.5uM) before imaging. TIRF images were acquired using a TIRF video microscope (Nikon) equipped with CFI Apo TIRF 100A-NA 1.49 oil objective and an EMC CD camera (Photometrics HQ2). The surface CAV1 puncta were quantified using the ImageJ (NIH) software.

### Live-cell surface biotin labeling to assess the cell surface Orai1 levels

Cell monolayers were washed with ice-cold PBS, and incubated in the same buffer containing sulfo-NHS-LC-biotin (#A39257, Thermo Fisher) for 30 min at 4°C. The cells were washed in PBS and their lysates in TX-100 lysis buffer subjected to anti-Orai1 immunoprecipitation followed by blotting with Streptavidin-Horseradish Peroxidase (HRP) Conjugate (cat. # SA10001) and enhanced chemiluminescence detection.

### Calcium flux assays

Cells were seeded in 35-mm glass-bottom dishes (cat. #FD35-100, WPI Inc) and loaded with Fluo4-AM in modified Tyrode’s solution (2 mM calcium chloride, 1 mM magnesium chloride, 137 mM sodium chloride, 2.7 mM potassium chloride, 12 mM sodium bicarbonate, 0.2 mM sodium dihydrogen phosphate, 5.5 mM glucose, pH 7.4) for 1 hour. After washing with calcium-free Tyrode’s solution, live cells were imaged under a confocal microscope (LSM710; Carl Zeiss), with fluorescence excitation at 488 nm and emission at 490– 540 nm. To initiate the release of intracellular Ca2+ stores, cells were stimulated with 2.5 μM thapsigargin in the absence of extracellular Ca^2+^. Once the signals approached the baseline, calcium chloride was added to 2 mM final concentration to record the SOCE (59). Data are presented as fold change in fluorescence emission relative to baseline.

### Patient population and tissue microarrays

Tissue microarrays (TMAs) corresponding to a well-annotated 971 breast cancer patient cohort at the University of Nottingham Hospital Breast Unit were analyzed by IHC staining with a previously described anti-rabbit EHD2 antibody (19) that was further validated (See Supplementary Fig. S1). Of the whole series (840 cases), 759 were informative. Both cytoplasmic and nuclear EHD2 signals were recorded. For statistical analysis, the expression was dichotomized using cutoff points that were selected based on histogram distribution using the median and X-tile software as follows: a H-score of zero for nuclear EHD2 and H-score of 50 for cytoplasmic EHD2 expression. Statistical analysis was performed using the SPSS IBM 22 statistical software (SPSS Inc., Chicago, IL, USA). The relationship between nuclear and cytoplasmic EHD2 expression and different clinic pathological variables was assessed using Chi square-test. Survival curves were obtained using Kaplan–Meier method for outcome estimation and significance determined using the log-rank test. Two-tailed P-values less than 0.05 were considered significant. Multivariate analysis was performed using the Cox hazard analysis.

### Prognostic analysis and gene targeted correlation analysis of *EHD2, CAV1* and *CAV2* mRNAs

The Kaplan–Meier plotter was used to evaluate the prognosis value of *EHD2*, *CAV1* and *CAV2* mRNA expressions alone and in combination (60). To analyze the survival probability alone and in combinations of *EHD2*, *CAV1* and *CAV2* mRNA, the patient cohorts were split on the basis of trichotomization (T1 vs T3). The *EHD2* mRNA (probe set 221870_at), *CAV1* mRNA (probe set 212097_at), and *CAV2* mRNA (probe set 203323_at) were entered into the KM Plotter patient cohort basal-like (PAM50 subtype) patient cohort (n=953) and the relapse free survival (RFS) was determined. The mean expression of *EHD2, CAV1* and *CAV2* were used to perform survival analysis of high *EHD2, CAV1* and *CAV2* vs. low *EHD2, CAV1* and *CAV2*. The hazard ratio (HR) with 95% confidence and log rank p-values were obtained from KM plotter. Gene correlation targeted analysis was performed to assess the correlation between *EHD2*, *CAV1* and CAV2 mRNA expression in basal-like (PAM50 subtype) breast cancer patients (n=783) and TNBC (IHC) cohort (n=293) using bc-GenExMiner v4.5 platforms. TCGA and SCAN-B RNAseq dataset were used. Correlation heatmap, correlation plots and Pearson’s correlation coefficients computation were performed using bc-GenExMiner v4.5.

### Statistical analysis

Statistical analysis of in vitro data was performed by comparing groups using unpaired student’s t test. In vivo tumorigenesis and metastasis data were analyzed using two-way ANOVA. A p value of <0.05 was considered significant.

### Human and animal subjects

The Ethics Committee of University of Nottingham approved the use of human tissues. All mouse xenograft and treatment studies were pre-approved by the UNMC Institutional Animal Care and Use Committee (IACUC) and conducted strictly according to the pre-approved procedures, in compliance with Federal and State guidelines.

## Acknowledgements

We thank Drs. Anjana Rao for CFP-STIM1, Ari Helenius for Cav1-EGFP and Ian Frew for the pMule kit (used to assemble the tdTomato/luciferase) through Addgene. This research was funded by grants from DOD (W81XWH-17-1-0616 and W81XWH-20-1-0058 to HB and W81XWH-20-1-0546 to VB) and NIH (R21CA241055 and R03CA253193 to VB), and by Fred & Pamela Buffett Cancer Center (pilot grants to HB & VB) and the Raphael Bonita Memorial Fund. The UNMC core facilities are supported by the NCI Cancer Center Support Grant (P30CA036727) to Fred & Pamela Buffett Cancer Center and the Nebraska Research Initiative. TAB, AMB and SC received University of Nebraska Medical Center Graduate Student Fellowships.

## Supplementary Figure Legends

**Supplementary Figure 1S. EHD2 is expressed in basal-like mammary epithelial cells and tumor cell lines.** (A) Immunoblot (left) and immunofluorescence (right) analysis of wildtype (EHD2+/+) and EHD2-null (EHD2-/-) mouse mammary gland to validate the specific reactivity of anti-EHD2 antibody used in this study. (B) Immunofluorescence analysis of EHD2 expression in basal vs. luminal epithelial cells of normal mouse mammary gland. Top panel, EHD2 (red) co-staining with basal cell marker alpha smooth muscle actin (SMA; green); Bottom panel, EHD2 (red) co-staining with luminal cell marker cytokeratin 8 (CK8; green). Nuclei are stained with DAPI (blue). Scale bars, 20 μm. (C) Confirmation of the basal epithelial cell-selective EHD2 expression in mouse mammary gland by immunohistochemical staining. Scale bars, 100 μm. (D) Predominant basal epithelial cell expression of EHD2 revealed by immunoblot analysis of Matrigel-grown organoids derived from FACS-sorted EPCAM-low/CD29-high (basal) vs. EPCAM-high/CD29-low (luminal) mouse mammary epithelial cell populations.

**Supplementary Figure S2. High cytoplasmic EHD2 correlated with shorter survival of breast cancer patients.** (A) Immunoblot analysis of EHD2 expression in non-tumorigenic immortal basal-like (76Ntert, MCF10a), and luminal A (ER+/PR+), luminal B (ER+/PR+, ErbB2+), ErbB2+, and Triple-negative (TN) breast cancer cell lines. (B) Immunofluorescence microscopy analysis of selected cell lines from A to further validate EHD2 (Red) expression pattern, showing predominant cytoplasmic and membrane localization D2. DAPI (blue) marks the nuclei. Scale bar, 50 μm. (C) EHD2 mRNA expression in breast cancer cell lines corresponding to major molecular subtypes as described in (23). (D) EHD2 mRNA expression in breast cancer cell lines corresponding to TNBC subtypes as described in (24). In C and D, The CCLE mRNA expression data is obtained as follows (per the CCLE site): RNASeq files are aligned with STAR and quantified with RSEM, and then TPM normalized. Reported values are Log2 (TPM+1); TPM, transcripts per million. The dotted line represents the median expression value of EHD2 among all (N=63) BC cell lines. The black and red asterisks (*) in C indicate cell lines that we show as negative or positive for EHD2 expression by western blotting or immunofluorescence microscopy (Fig. S2A and S2B). (E) Kaplan-Meier survival curve for Breast Cancer Specific Survival (BCSS) probability in all tumors scored for cytoplasmic and nuclear positive (purple), cytoplasmic and nuclear negative (blue), cytoplasmic positive and nuclear negative (green), cytoplasmic negative and nuclear positive (yellow) EHD2 expression. N= 275 for cytoplasmic and nuclear negative, 183 for cytoplasmic positive and nuclear negative, 76 for cytoplasmic negative and nuclear positive and 207 for cytoplasmic and nuclear positive.

**Supplementary Figure S3. Knockdown of EHD2 impaired the tumorigeneses and metastases in vivo.** Primary tumor xenograft tumorigenesis and lung metastasis of MDA-MB-231 cells expressing control or EHD2 shRNA. Representative primary tumors and lungs are shown on left. Table on right shows the numbers of mice developing identifiable primary tumors and lung metastases.

**Supplementary Figure S4. EHD1/4 expression is unchanged in EHD2 knockout TNBC cell lines.** (A) Immunoblot analysis of EHD1/4 expression in EHD2 WT and KO MDA-MB-231, Hs578t and BT549 breast cancer cell lines. (B) Densitometric quantification of EHD1/4 expression levels from three independent experiments; ns, not significant (Student’s t test).

**Supplementary Figure S5. Loss of EHD2 decreased ontogenesis traits in TNBC cells.** (A) Representative images of cell migration in control and EHD2 KO TNBC cells. (B) Representative images of cell invasion in control and EHD2 KO TNBC cells. (C) Representative images of extracellular matrix degradation in control and EHD2 KO TNBC cells.

**Supplementary Figure S6. EHD2 and Caveolin-1/2 are correlated in breast cancers patients.** (A) Pearson’s pairwise correlation heatmap analysis of the expression of targeted genes (*EHD2, CAV1* and *CAV2*) for the TNBC cohort based on IHC. Analysis used TCGA and SCAN-B RNAseq dataset of bc-GenExMiner v4.5 platform. (B) KM plotter analysis of upper vs. lower quartile survival curves display poor relapse-free survival (RFS) associated with EHD2, CAV1 and CAV2 overexpression in basal-like breast cancer (based on PAM50 subtype) cohorts of TCGA, GEO and GEA datasets (KM plotter). The probe sets used were: EHD2 (221870_at), CAV1 (212097_at) and CAV2 (203323_at). Similar analysis on all breast cancer samples in the cohort found no survival differences (lower panel).

**Supplementary Figure S7. Loss of EHD2 decreased SOCE in TNBC cells.** Enhancement of reporter fluorescence measured upon thapsigargin (2.5 μM) and Ca^2+^ (2mM), immunofluorescent images of SOCE before and after Ca^2+^ addition from Control and EHD2 KO Hs578T cells engineered with the cytoplasmic calcium sensor GCaMP6s (A) or plasma membrane-localized calcium sensor GCaMP6s-CAAX (B).

**Supplementary Figure S8. Validation of ORAI1 antibody in triple negative breast cancer cell lines.** MDA-MB-231 and BT-549 were transfected with either control siRNA or Orai1 siRNA for 72h. (A) Whole cell lysates were subjected to immunoblotting with anti-Orai1 (cat. # O8264, Sigma). Orai1appears as a smear. (B) Total RNA was used for Orai1 qPCR to establish successful and specific knockdown of Orai1 mRNA expression in Orai1 siRNA transfected cells.

