## Supplemental Tables for "EHD2 overexpression promotes tumorigenesis and metastasis in triple-negative breast cancer by regulating store-operated calcium entry"

**Supplementary Table**

**Supplementary Table S1. (A) Validation summary of EHD2 staining from all patients.**

|  | Statistics | |
| --- | --- | --- |
|  | (Nuclear) N EHD2 | (Cytoplasmic) C EHD2 |
| Valid | 756 | 759 |
| Mean | 40.99 | 51.36 |
| Median | 0 | 10 |
| Std. Deviation | 75.27 | 74.59 |

**Supplementary Table S1. (B) Associations between EHD2 nuclear and cytoplasmic expression and clinical, pathological, and biological characteristics in the complete patient series.**

|  | **EHD2 Nuclear Expression** | | |  | **EHD2 Cytoplasmic Expression** | | |
| --- | --- | --- | --- | --- | --- | --- | --- |
|  | Negative | Positive | *p*-value |  | Negative | Positive | *p*-value |
|  | (=0) | (>0) | (χ^2^) |  | (<50) | (≥50) | (χ^2^) |
| **Age** |  |  |  |  |  |  |  |
| <50 | 163(36) | 100(33) | 0.449 |  | 123(34) | 140(35) | 0.796 |
| ≥50 | 288(64) | 199(67) |  |  | 234(66) | 256(65) |  |
| **Menopausal Status** |  |  |  |  |  |  |  |
| Pre- | 180(40) | 111(37) | 0.539 |  | 143(40) | 149(37) | 0.463 |
| Post- | 276(61) | 187(63) |  |  | 215(60) | 250(63) |  |
| **Tumour Size (cm)** |  |  |  |  |  |  |  |
| < 2.0 | 185(41) | 165(56) | **<0.001** |  | 172(49) | 179(46) | 0.422 |
| ≥2.0 | 262(59) | 129(44) |  |  | 181(51) | 212(54) |  |
| **Stage** |  |  |  |  |  |  |  |
| 1 | 272(60) | 185(63) | 0.303 |  | 221(63) | 238(61) | 0.847 |
| 2 | 140(31) | 92(31) |  |  | 107(30) | 126(32) |  |
| 3 | 38(9) | 16(6) |  |  | 25(7) | 29(7) |  |
| **Grade** |  |  |  |  |  |  |  |
| 1 | 45(10) | 62(21) | **<0.001** |  | 58(17) | 49(12) | **0.012** |
| 2 | 119(27) | 103(35) |  |  | 118(33) | 105(27) |  |
| 3 | 285(64) | 128(44) |  |  | 176(50) | 239(61) |  |
| **Mitosis** |  |  |  |  |  |  |  |
| 1 | 101(23) | 111(39) | **<0.001** |  | 120(35) | 92(24) | **0.006** |
| 2 | 81(18) | 59(21) |  |  | 63(18) | 78(20) |  |
| 3 | 261(59) | 114(40) |  |  | 163(47) | 214(56) |  |
| **Vascular Invasion** |  |  |  |  |  |  |  |
| Probable/Negative | 263(59) | 208(71) | **0.001** |  | 225(64) | 248(64) | 0.925 |
| Definite | 183(41) | 85(29) |  |  | 127(36) | 142(36) |  |
| **ER** |  |  |  |  |  |  |  |
| Negative | 145(32) | 67(23) | **0.005** |  | 92(26) | 122(31) | 0.151 |
| Positive | 307(68) | 229(77) |  |  | 262(74) | 275(69) |  |
| **PgR** |  |  |  |  |  |  |  |
| Negative | 209(48) | 101(35) | **0.001** |  | 142(42) | 170(45) | 0.419 |
| Positive | 226(52) | 185(65) |  |  | 200(58) | 212(55) |  |
| **HER2 Status** |  |  |  |  |  |  |  |
| Negative | 361(83) | 244(86) | 0.294 |  | 304(87) | 304(82) | 0.039 |
| Positive | 74(17) | 40(14) |  |  | 45(13) | 69(18) |  |
| **AR** |  |  |  |  |  |  |  |
| Negative | 195(49) | 88(34) | **<0.001** |  | 146(46) | 139(41) | 0.196 |
| Positive | 206(51) | 170(66) |  |  | 174(54) | 203(59) |  |
| **Her3** |  |  |  |  |  |  |  |
| Negative | 29(7) | 25(10) | 0.204 |  | 28(9) | 26(7) | 0.555 |
| Positive | 386(93) | 232(90) |  |  | 296(91) | 325(93) |  |
| **Her4** |  |  |  |  |  |  |  |
| Negative | 42(10) | 36(13) | 0.177 |  | 41(12) | 37(10) | 0.391 |
| Positive | 394(90) | 244(87) |  |  | 304(88) | 337(90) |  |
| **Ck5** |  |  |  |  |  |  |  |
| Negative | 278(79) | 192(88) | **0.008** |  | 240(87) | 230(79) | **0.011** |
| Positive | 72(21) | 26(12) |  |  | 37(13) | 63(21) |  |
| **Ck7/8** |  |  |  |  |  |  |  |
| Negative | 10(2) | 5(2) | 0.622 |  | 5(1) | 11(3) |  |
| Positive | 425(98) | 279(98) |  |  | 340(99) | 366(97) | 0.181 |
| **Ck14** |  |  |  |  |  |  |  |
| Negative | 378(91) | 239(89) | 0.444 |  | 297(90) | 322(90) | 0.925 |
| Positive | 39(9) | 30(11) |  |  | 34(10) | 36(10) |  |
| **Ck17** |  |  |  |  |  |  |  |
| Negative | 269(82) | 170(89) | **0.039** |  | 221(87) | 219(82) | 0.116 |
| Positive | 58(18) | 21(11) |  |  | 33(13) | 48(18) |  |
| **HER1** |  |  |  |  |  |  |  |
| Negative | 338(77) | 227(79) | 0.451 |  | 280(81) | 287(75) | 0.06 |
| Positive | 101(23) | 59(21) |  |  | 66(19) | 95(25) |  |
| **Mucin1** |  |  |  |  |  |  |  |
| Negative | 50(12) | 15(6) | **0.005** |  | 28(9) | 39(11) | 0.367 |
| Positive | 355(88) | 245(94) |  |  | 286(91) | 315(89) |  |
| **P53** |  |  |  |  |  |  |  |
| Negative | 292(68 | 207(74) | 0.073 |  | 245(72) | 255(68) | 0.255 |
| Positive | 138(32) | 72(26) |  |  | 94(28) | 118(32) |  |
| **Mdm2** |  |  |  |  |  |  |  |
| Negative | 264(82) | 152(71) | **0.003** |  | 199(76) | 219(78) | 0.531 |
| Positive | 60(19) | 63(29) |  |  | 63(24) | 61(22) |  |
| **Ki67** |  |  |  |  |  |  |  |
| Negative | 99(29) | 106(45) | **<0.001** |  | 105(38) | 101(33) | 0.157 |
| Positive | 245(71) | 132(56) |  |  | 170(62) | 209(67) |  |
| **ADA3** |  |  |  |  |  |  |  |
| Negative | 201(53) | 93(40) | **0.002** |  | 149(52) | 147(44) | 0.058 |
| Positive | 181(47) | 141(60) |  |  | 138(48) | 185(56) |  |
| **TN Status** |  |  |  |  |  |  |  |
| Non-TN | 338(77) | 254(87) | **0.001** |  | 288(82) | 305(79) | 0.293 |
| TN | 102(23) | 38(13) |  |  | 62(18) | 80(21) |  |
