## Supplemental Figures for "EHD2 overexpression promotes tumorigenesis and metastasis in triple-negative breast cancer by regulating store-operated calcium entry"

Supplementary Fig. S1

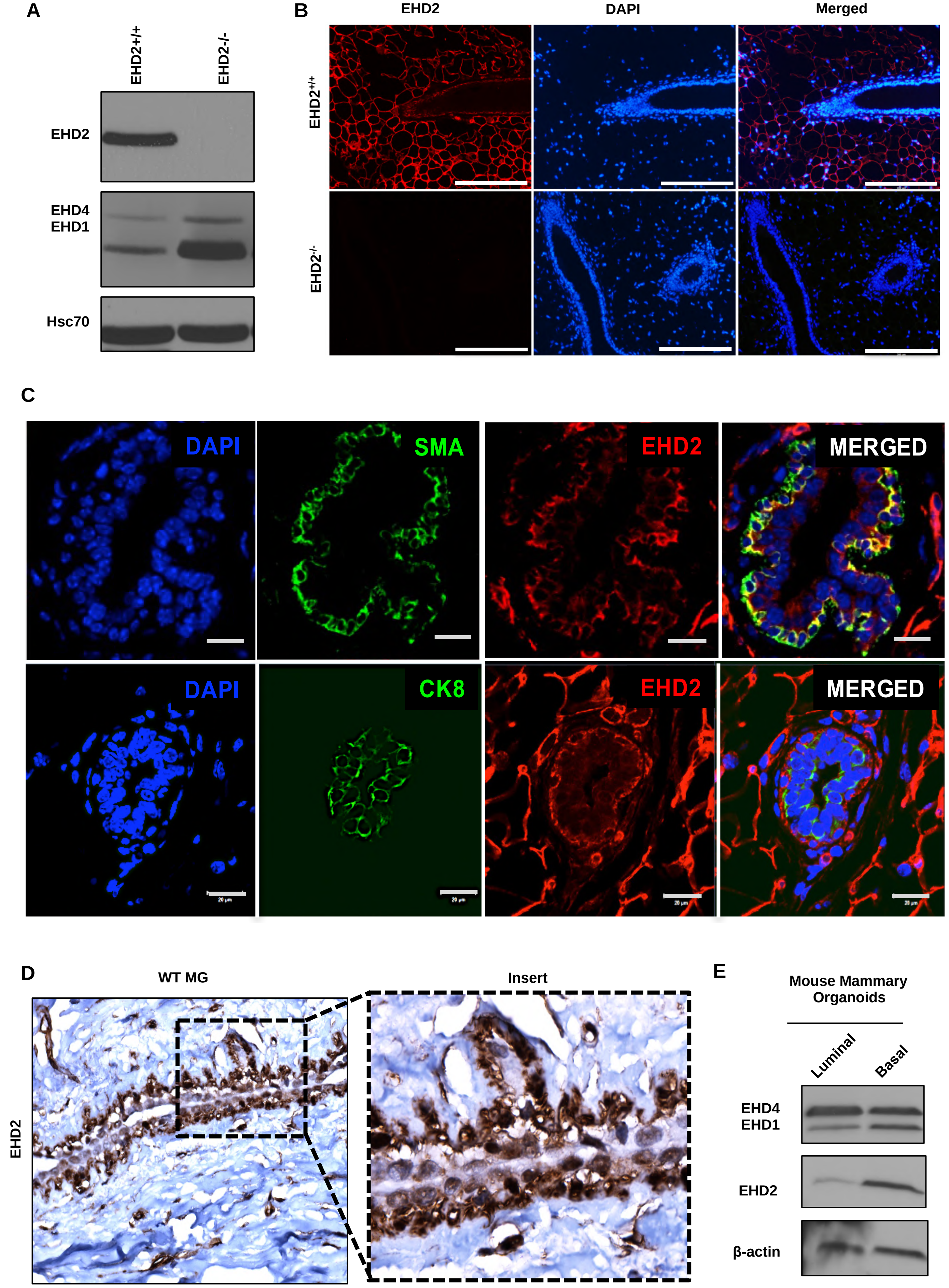

**Supplementary Fig. S2**

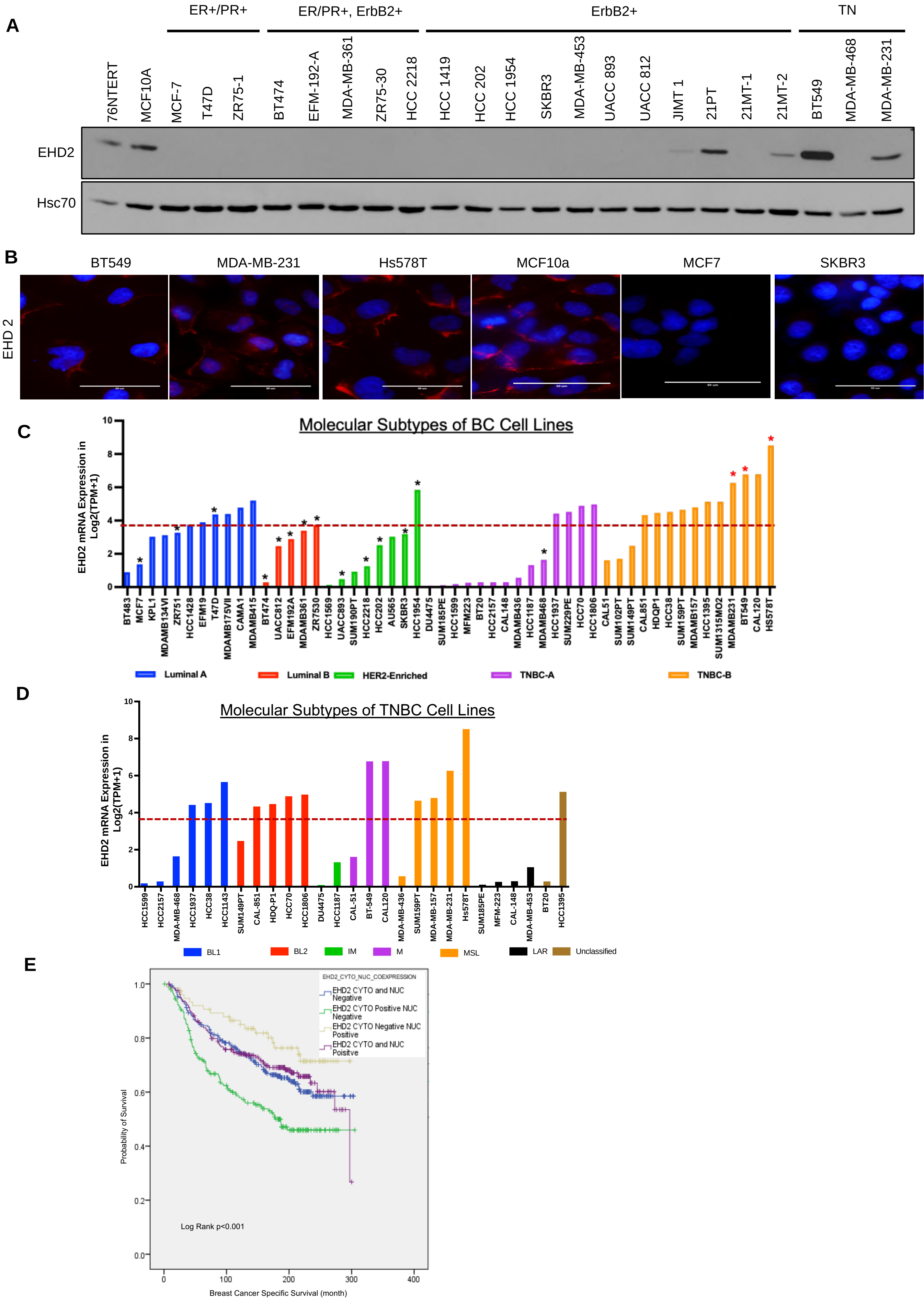

### Supplementary Fig. S3

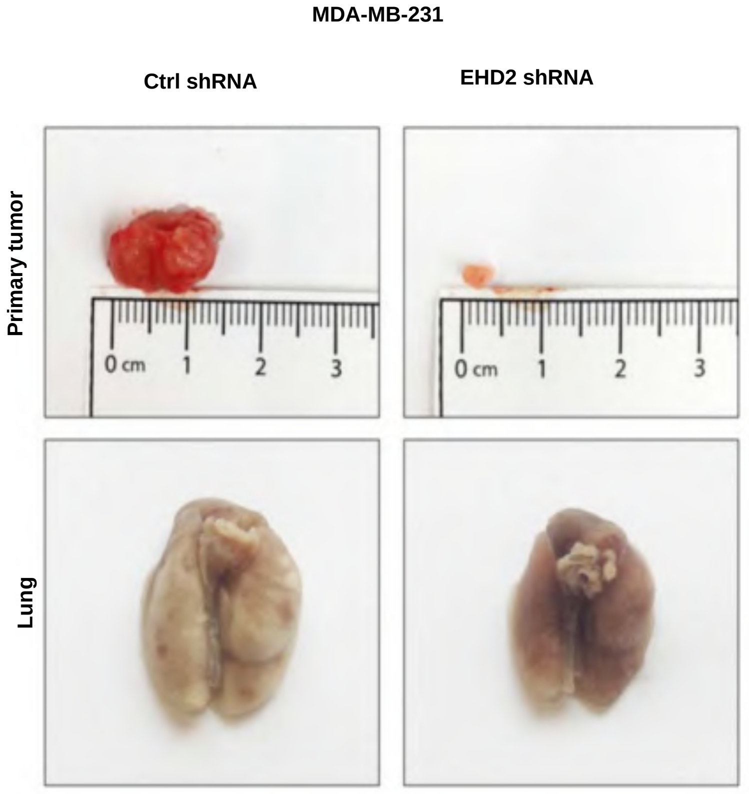


| MDA-MB-231 | Ctrl shRNA | EHD2 shRNA |
| --- | --- | --- |
| Tumors Formed | 6/6 | 4/6 |
| Lung Metastases | 5/6 | 1/4 |

**Supplementary Fig. S4**

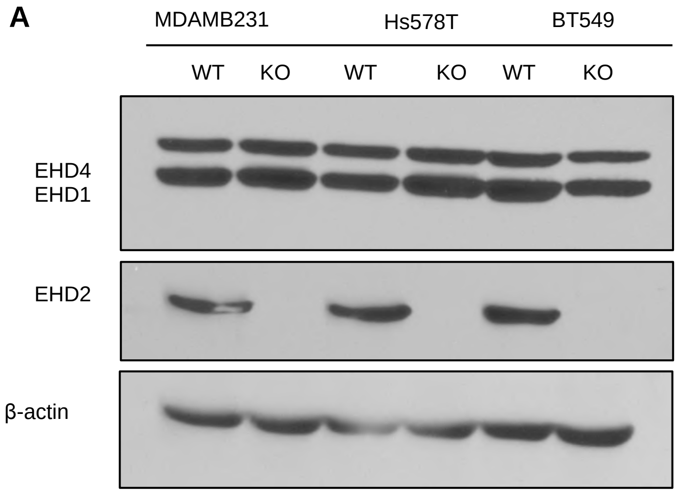


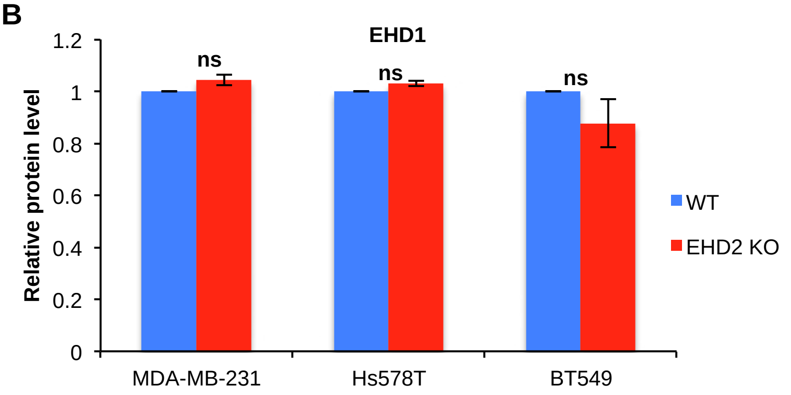


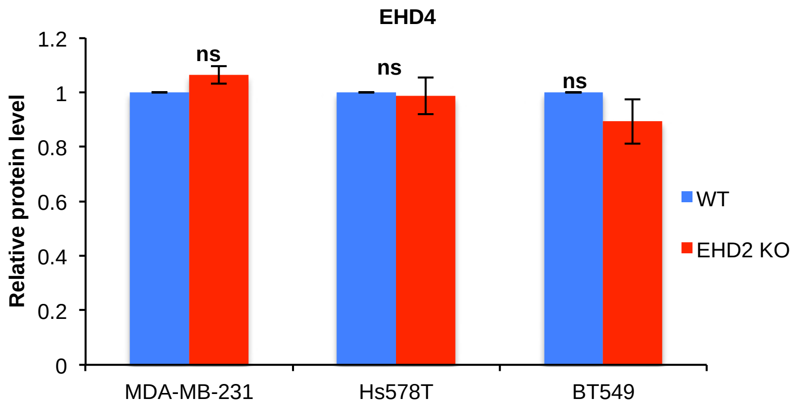

Supplementary Fig. S5

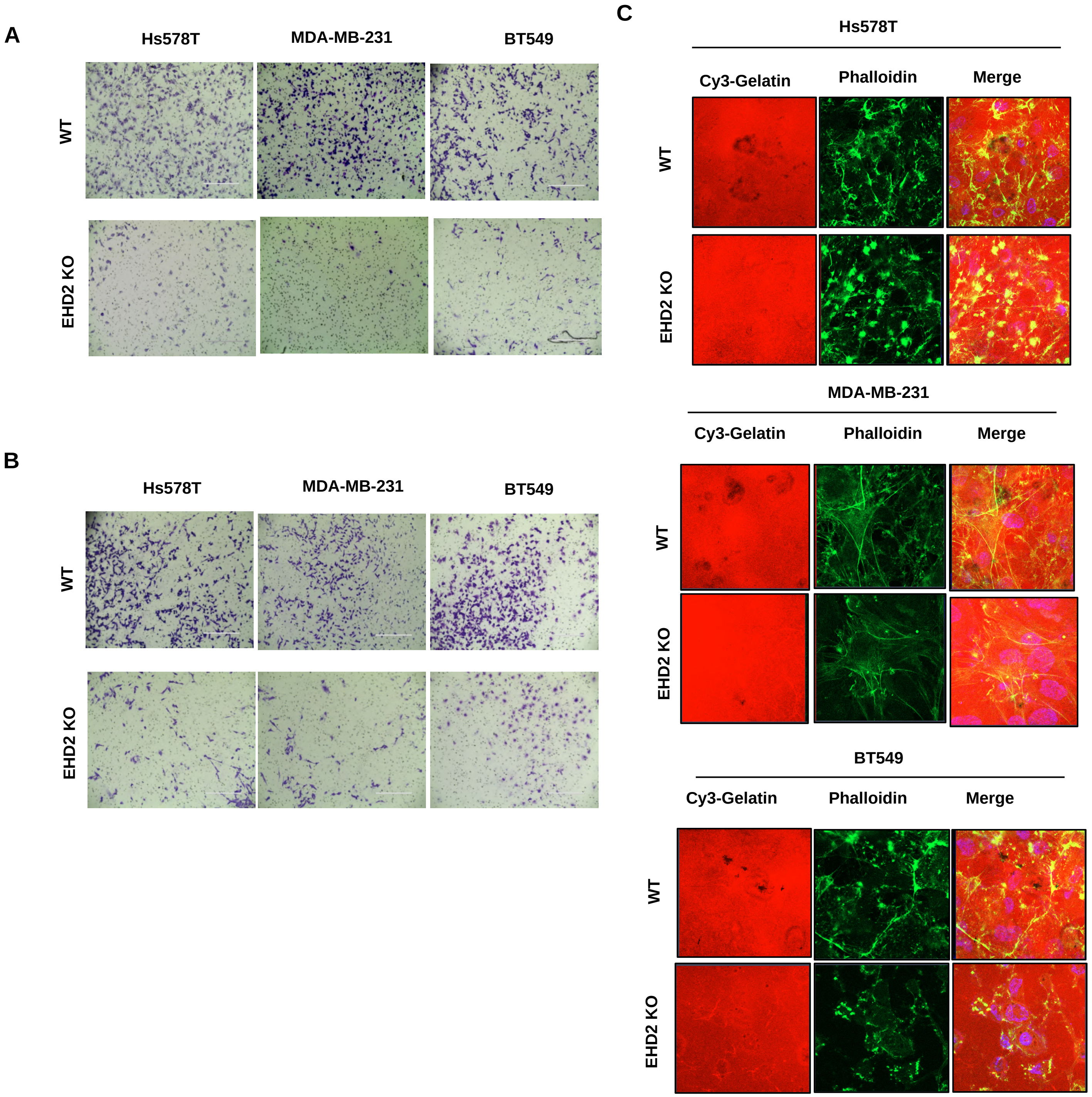

### Supplementary Fig. S6

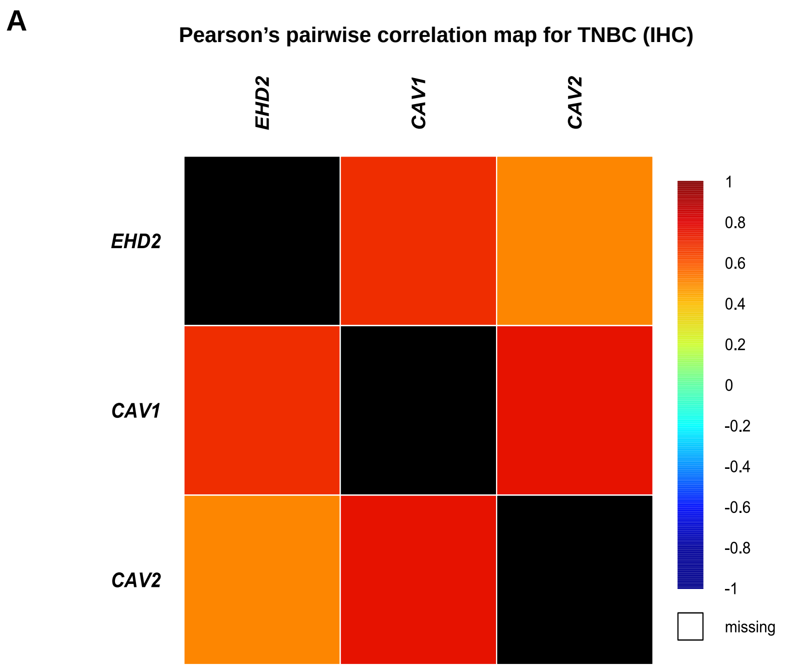


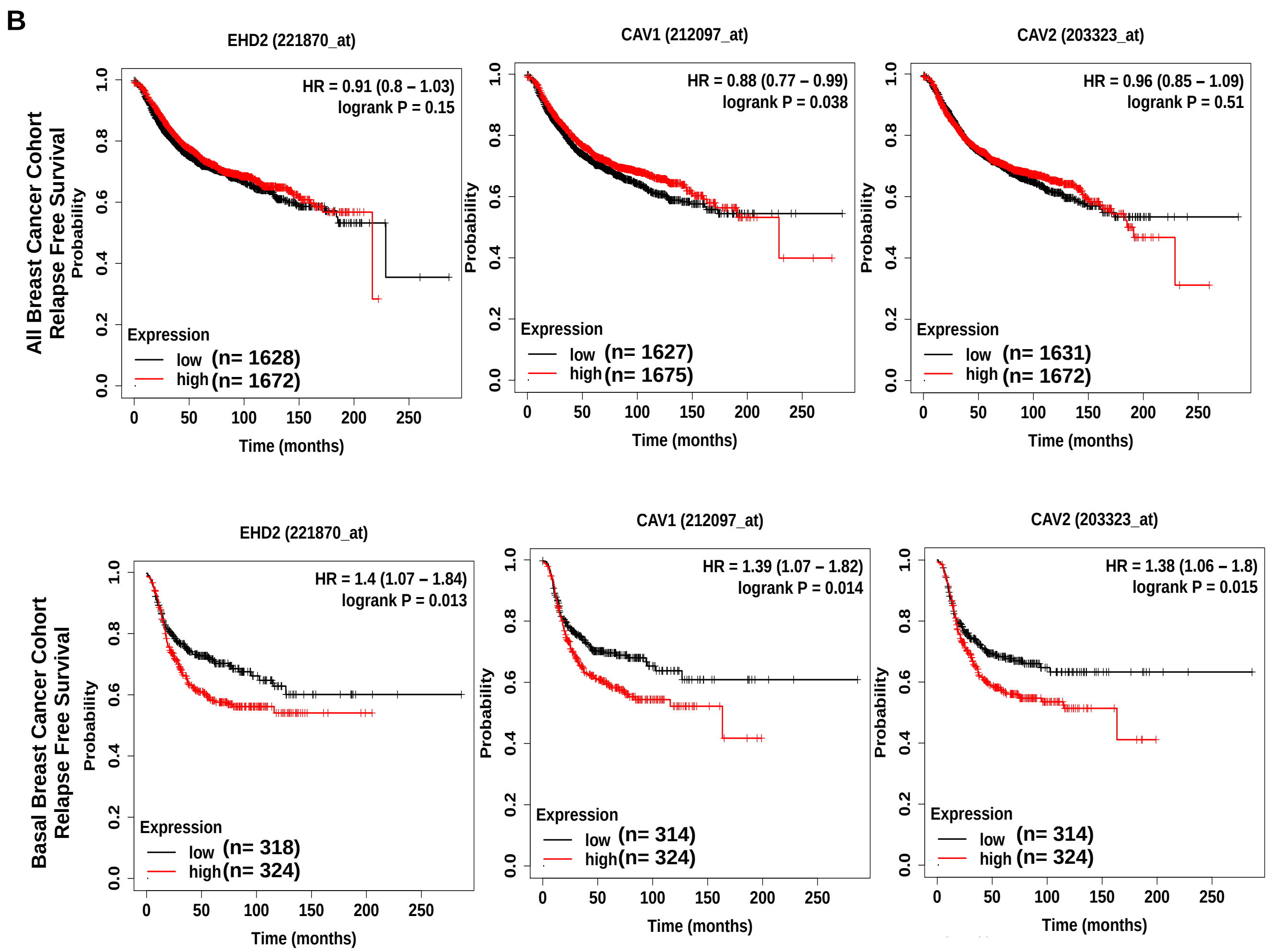

### Supplementary Fig. S7

A

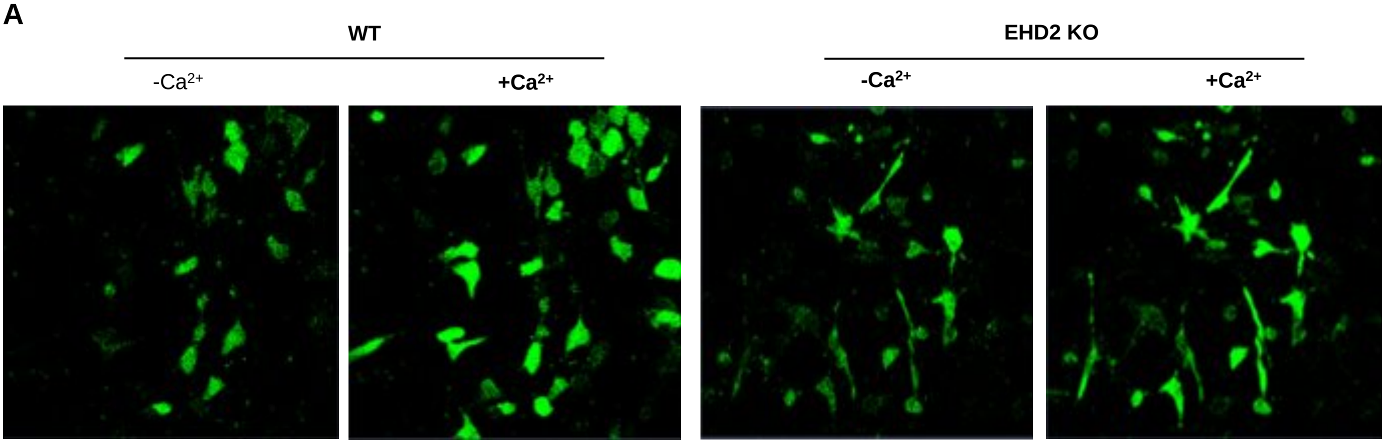


B

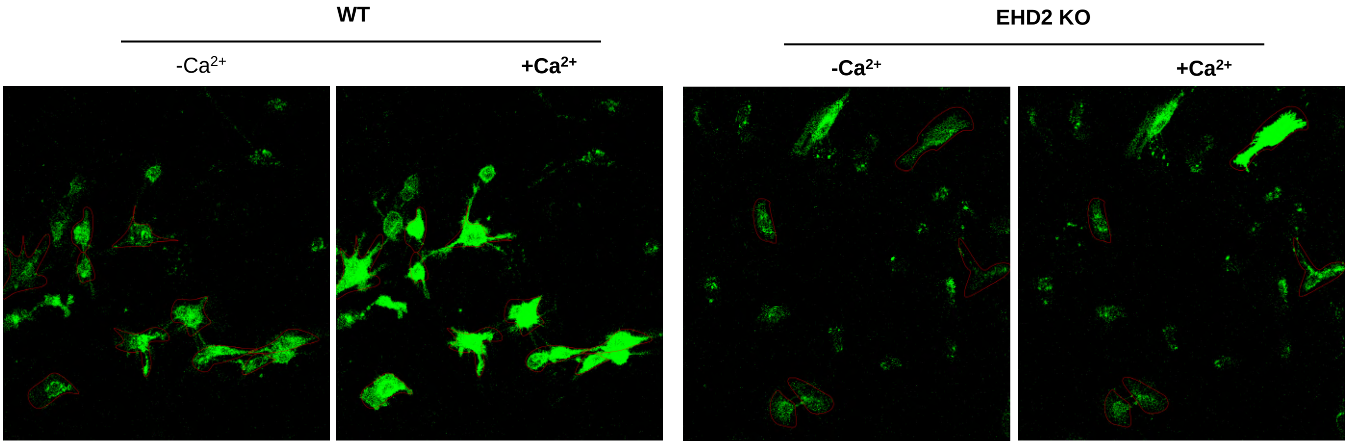

**Supplementary Fig. S8**

**A**

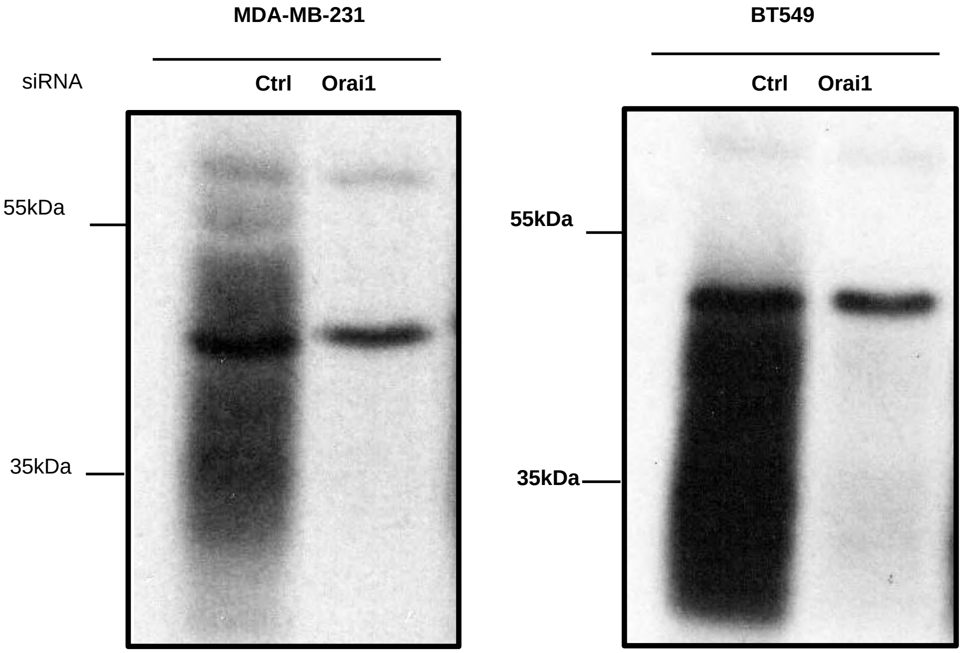


**B**

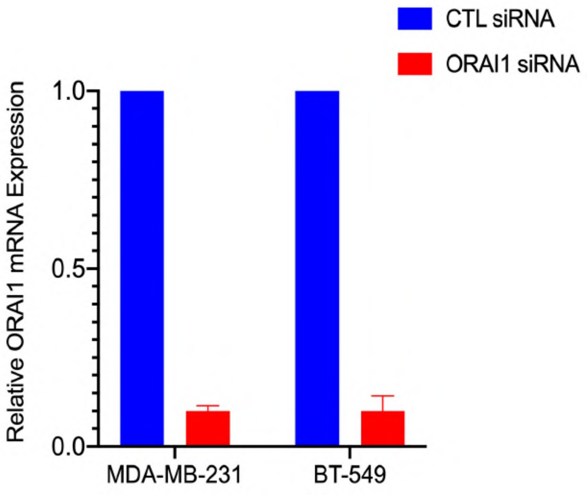
